# Positional cues underlie cell fate specification during branching morphogenesis of the embryonic mammary epithelium

**DOI:** 10.1101/2022.08.30.505826

**Authors:** Claudia Carabaña, Wenjie Sun, Meghan Perkins, Varun Kapoor, Robin Journot, Fatima Hartani, Marisa M. Faraldo, Bethan Lloyd-Lewis, Silvia Fre

## Abstract

How cells coordinate morphogenetic cues and fate specification during development is a fundamental question at the basis of tissue formation. Lineage tracing studies have demonstrated that many stratified epithelia, including the mammary gland, first arise from multipotent stem cells, which are progressively replaced by distinct pools of unipotent progenitors that maintain tissue homeostasis postnatally. The lack of specific markers for early fate specification in the mammary gland has prevented the delineation of the features and spatial localization of lineage-committed progenitors that co-exist with multipotent stem cells (MaSCs) during tissue development. Here, using single-cell RNA-sequencing across 4 stages of embryonic development, we reconstructed the differentiation trajectories of multipotent mammary stem cells towards basal and luminal fate. Our data revealed that MaSCs can already be resolved into distinct populations exhibiting lineage commitment at the time coinciding with the first sprouting events of mammary branching morphogenesis (E15.5). By visualizing gene expression across our developmental atlas, we provide novel molecular markers for committed and multipotent MaSCs, and define their spatial distribution within the developing tissue. Furthermore, we show that the mammary embryonic mesenchyme is composed of two spatially-restricted cell populations, representing the sub-epithelial and dermal mesenchyme. Mechanistically, we explored the communication between different subsets of mesenchymal and epithelial cells, using time-lapse analysis of mammary embryonic explant cultures, and reveal that mesenchymal-produced FGF10 accelerates embryonic mammary branching morphogenesis without affecting cell proliferation. Altogether, our data elucidate the spatiotemporal signals underlying lineage specification of multipotent mammary stem cells and uncover the paracrine interactions between epithelial and mesenchymal cells that guide mammary branching morphogenesis.

## Introduction

To generate functional organs, cell fate acquisition and multicellular morphogenetic events must be tightly coordinated. Accordingly, lineage commitment encompasses a progressive differentiation process dictated by transcriptional and mechanical changes to drive the formation of specialist tissues of complex shapes and function (Chan et al., 2017). The development of the branched mammary gland (MG) is a case in point, being initially formed from multipotent embryonic mammary stem cells (MaSCs) which reorganize through individual and collective movements during branching morphogenesis until committing to specific luminal and basal lineages at birth. Subsequently, unipotent progenitors drive adult homeostasis (Blaas et al., 2016; Davis et al., 2016; Lilja et al., 2018; Lloyd-Lewis et al., 2018; Prater et al., 2014; Scheele et al., 2017; van Amerongen et al., 2012; van Keymeulen et al., 2011; Wuidart et al., 2016, 2018). The embryonic mammary gland therefore represents a powerful tissue paradigm to study the integration of stem cell fate specification with tissue morphogenesis.

Mouse mammary gland development begins at embryonic day (E) 10 with the formation of bilateral milk lines, followed by the asynchronous appearance of five pairs of epithelial placodes positioned symmetrically at each side of the embryo. By E13, these placodes invaginate into the underlying mesenchyme to give rise to mammary buds. At around E15.5, the epithelium undergoes the first sprouting event to invade the underlying fat pad precursor, triggering branching morphogenesis and the formation of a small rudimentary ductal tree by birth (reviewed in (Watson & Khaled, 2020)). The mammary ductal network is composed of a bilayered epithelium comprising two main cell types: an outer layer of myoepithelial or basal cells (BCs) adjacent to the basement membrane and an inner layer of polarized luminal cells (LCs), facing the ductal lumen, that encompass hormone receptor (namely Estrogen (ERα) and Progesterone (PR) receptors) expressing and non-expressing subpopulations.

We have recently shown that the lineage bias of MaSCs occurs progressively within a narrow developmental window around embryonic day E15.5, a surprisingly early time in mammogenesis (Lilja et al., 2018). Strikingly, this bias towards luminal and basal cell fates coincides with the remarkable epithelial remodeling that occurs during the first embryonic mammary branching event. Yet, the precise timing of lineage specification during this crucial stage of mammary gland morphogenesis remains unclear, hampered by the lack of specific markers for early fate specification.

It is now well-established that cell-fate-specific changes in gene expression can modify the properties of a growing tissue and affect its morphogenesis and patterning. In the mammary epithelium, recent studies performed single-cell RNA sequencing (scRNA-seq) analysis at distinct stages of mammary embryonic development and proposed a model whereby multipotent MaSCs drive the earliest stages of mammogenesis. These studies identified subsets of embryonic mammary cells characterized by ‘hybrid’ transcriptional signatures and harboring concomitant expression of luminal and basal genes (Giraddi et al., 2018; Wuidart et al., 2018). In contrast, alternative scRNA-seq studies suggested that only Mammary Epithelial Cells (MECs) with basal characteristics are present in the embryonic gland, and that these bipotent progenitors generate mammary luminal cells postnatally (Pal et al., 2021). Recent single nucleus Assay for Transposase Accessible Chromatin sequencing (snATAC-seq) analyses, however, revealed that MECs at E18.5 exhibit either a basal-like or luminal-like chromatin accessibility profile, suggesting the potential priming of these cells to a lineage-restricted state prior to birth (Chung et al., 2019).

Given these uncertainties, here we sought to further define the potency of mammary stem cells and the timing of fate acquisition with spatiotemporal resolution during embryonic mammary morphogenesis, by coupling single cell transcriptional mapping at different developmental timescales with *ex vivo* live imaging of mammary embryonic cell dynamics during branching morphogenesis. This enabled us to finely dissect the heterogeneity of the mammary gland epithelium throughout embryonic development and define the transcriptional programs orchestrating the lineage restriction of multipotent MaSCs to unipotent progenitors. Importantly, our integrative approach prospectively identified new markers for specific mammary cells, and provided fundamental insights into the resident mammary embryonic mesenchymal cells that direct branching morphogenesis.

## Results

### Lineage restriction is a progressive developmental process

How changes in mammary tissue architecture during morphogenesis translate into differential gene expression patterns that drive the lineage specification of individual cells during development remains unknown in many tissue contexts. To address this in the MG, we performed scRNA-seq analysis of mouse embryonic mammary tissues at four developmental times spanning mammary bud invagination (E13.5), initial sprouting events at the presumptive onset of lineage segregation (E14.5 and E15.5) (Lilja et al., 2018) and post-natal branching morphogenesis (at birth or Post-natal day 0, P0) (Figure 1A). At each timepoint, we micro-dissected mammary buds from female mouse embryos (pooling tissues from 7-12 embryos isolated from different pregnant dams) and isolated mammary epithelial (EpCAM^+^) and stromal (EpCAM^−^) cells by FACS for scRNA-seq using the 10x Chromium platform. Basal and luminal subpopulations are indistinguishable in embryonic mammary glands using the EpCAM and CD49f gating strategies routinely applied to adult tissues (Figure S1A).

**Figure 1.**
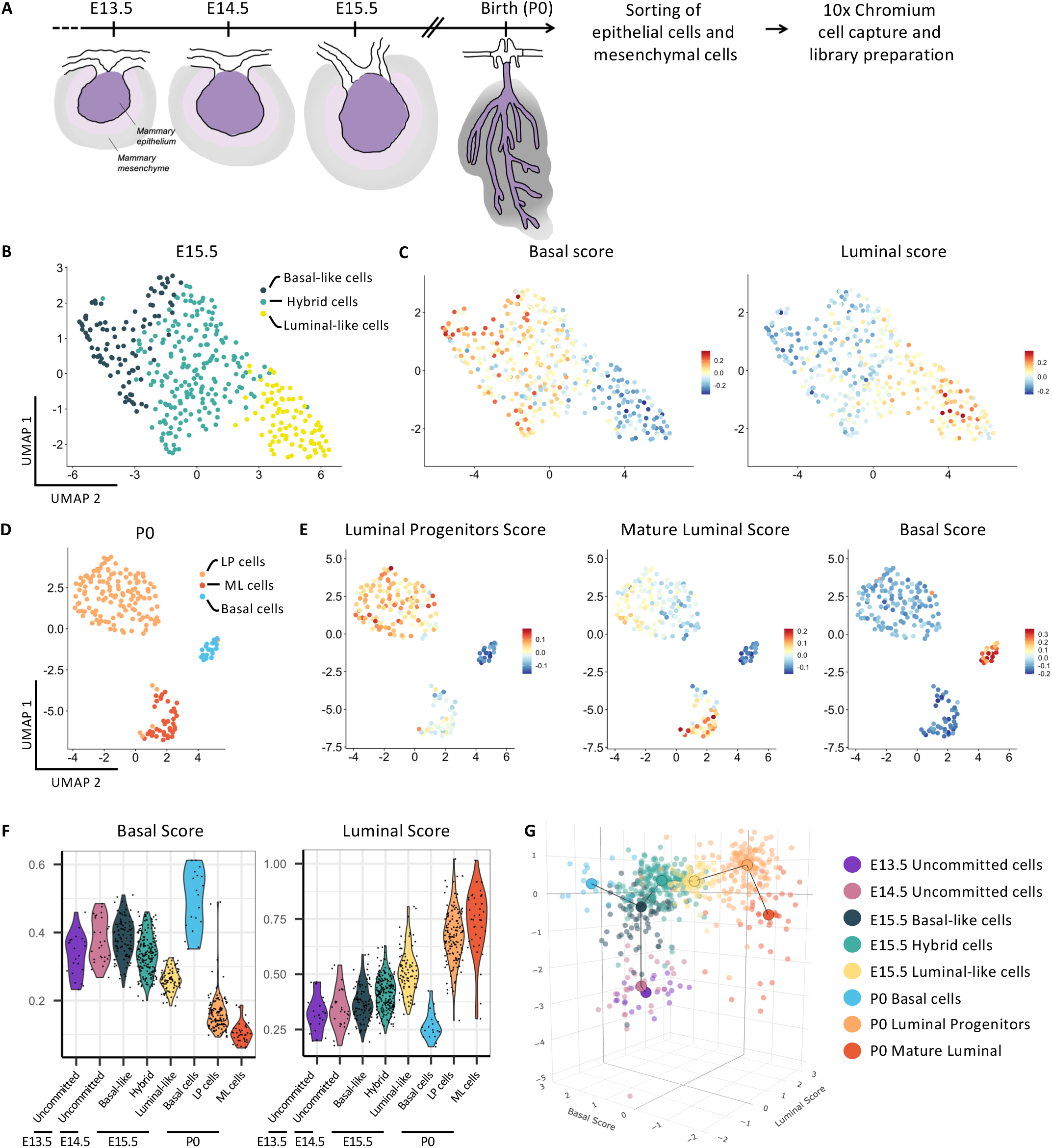
Developmental atlas of the transcriptional signatures and 3D trajectory analysis of luminal and basal differentiation of single mammary epithelial cells from E13.5 until birth. (A) Scheme showing the isolation and sequencing strategy of mammary embryonic cells at four developmental stages spanning embryonic MG development. (B) UMAP plot of embryonic MECs isolated at E15.5 after subset analysis of non-proliferative MG epithelial cells. Cells are color-coded by cluster. (C) UMAP plots from (B) color-coded according to the expression of the single-cell ID scores in MECs: basal score (left) and luminal score (right). (D) UMAP plot of MECs isolated at P0 after subset analysis of MG epithelial cells. (E) UMAP plots from (D) color-coded according to the expression of luminal progenitors (LP), mature luminal (ML) and basal cell (BC) scores. (F) Violin plots showing the expression levels of the basal and luminal scores in each cluster. (G) 3D trajectory of MECs from E13.5 at the origin of the mammary cellular hierarchy to P0 MECs positioned at the end of two divergent differentiation routes.

Using the Seurat R package (Stuart et al., 2019), unsupervised clustering of single cell expression data revealed distinct cell clusters at E13.5, E14.5, E15.5 and P0, respectively (Figure S1B), which were manually annotated by matching enriched gene sets with known markers of mammary epithelium, mesenchyme and skin cells. With the objective of mapping MECs undergoing lineage commitment early in embryogenesis, we removed contaminating skin cells (Figure S1B) and performed a sub-clustering analysis of epithelial populations at each developmental timepoint. A cluster composed of proliferative epithelial cells was identified at E15.5, based on a list of cell cycle related genes, which were omitted from further analysis (Figure S1C-D). While this analysis identified a single population of MECs at the early E13.5 and E14.5 developmental times, 3 transcriptionally distinct cell clusters were apparent at E15.5 and P0 (Figure 1B, 1D, S1B). The detection of 3 MECs clusters at E15.5 was surprising, as previous studies observed a single population around this developmental stage (Giraddi et al., 2018). To investigate this further, we calculated a single-cell ID score for “basal-like” and “luminal-like” cells based on previously published transcriptomic analyses of adult MECs (Kendrick et al., 2008). A higher single-cell ID score reflects increasing similarity to the reference cell type: adult basal or luminal cells. Interestingly, this analysis revealed that E15.5 MECs can already be resolved into 3 distinct groups: luminal-like cells, basal-like cells and a hybrid cell population co-expressing luminal and basal genes (Figure 1C, S1E). As expected, lineage markers commonly used to distinguish LCs (*Krt8, Krt18*) from BCs (*Krt5, Trp63*) in the postnatal mammary gland were co-expressed in all 3 MECs clusters at E15.5 (Figure S1F). Importantly, alongside established markers for adult BCs (*Lmo1, Pthlh, Cxcl14*) and LCs (*Anxa1, Ly6d*) (Kendrick et al., 2008), this analysis also identified genes that had not been previously ascribed to distinct mammary BC or LC populations.

By applying a computed ID score for each epithelial adult cell type (Kendrick et al., 2008) to the 3 transcriptionally distinct cell populations observed at P0 (Figure 1D), BCs (*Acta2*^*+*^, *Myh11*^*+*^), luminal progenitors (LP) (*Notch1*^*+*^, *Aldh1a3*^*+*^, *Lypd3*^*+*^) and mature luminal (ML) cells (*Prlr*^*+*^, *Cited1*^*+*^, *Esr1*^*+*^) could be clearly distinguished (Figure 1E). This corroborates our previous findings indicating that MECs are already committed to 3 distinct lineages at birth (Lilja et al., 2018). Moreover, these results are consistent with previous snATAC-seq analyses of the embryonic mammary gland, which also identified 3 separate clusters at E18.5 (Chung et al., 2019). Collectively, our data supports a model whereby mammary epithelial cell lineages are progressively being specified throughout development and are well segregated at birth.

We next ordered the cells along pseudo-temporal trajectories to infer the differentiation path of embryonic MECs towards luminal or basal fate. Since we observed that the 2^nd^ principal component of the PCA was highly correlated to the age of the embryos analyzed, we used it as a proxy for developmental stage (y-axis) and plotted it against the basal and luminal scores computed above (Figure 1C) on the x-axis (Kendrick et al., 2008) (Figure 1F-G, S1G). The resulting plot indicates, as predicted, that E13.5 mammary cells lie at the origin of the mammary cellular hierarchy, with E15.5 cell populations occupying intermediate positions and P0 MECs positioned at the end of two divergent trajectories, representing the binary cell fate choice between basal or luminal differentiation. Remarkably, we noticed that basal-like cells at E15.5 can either transition towards the P0 basal cluster, or to a hybrid cell state that will give rise to LCs (Figure 1G), suggesting that they might lie at the origin of both lineages.

Together, our temporal scRNA-seq atlas reveals the molecular changes associated with progressive lineage restriction and identifies subsets of MECs that are already biased towards basal or luminal cell fate at embryonic day E15.5. Thus, both committed (i.e. conceivably unipotent) and undifferentiated (putative multipotent) cells likely exist at this important developmental stage in mammogenesis, which coincides with the first morphogenetic events of mammary epithelial branching and duct elongation (Lilja et al., 2018).

### Luminal and basal progenitors are already spatially segregated at E15.5

We next sought to identify differentially expressed genes for each mammary epithelial cluster by examining their dynamic expression profile towards luminal or basal differentiation trajectories. While our compiled scRNA-seq atlas emphasized the vast cellular heterogeneity of the embryonic mammary epithelium, this extended analysis identified different patterns of expression along the process of basal (Figure 2A) or luminal (Figure 2B) differentiation throughout embryonic development (from E13.5 to P0).

**Figure 2.**
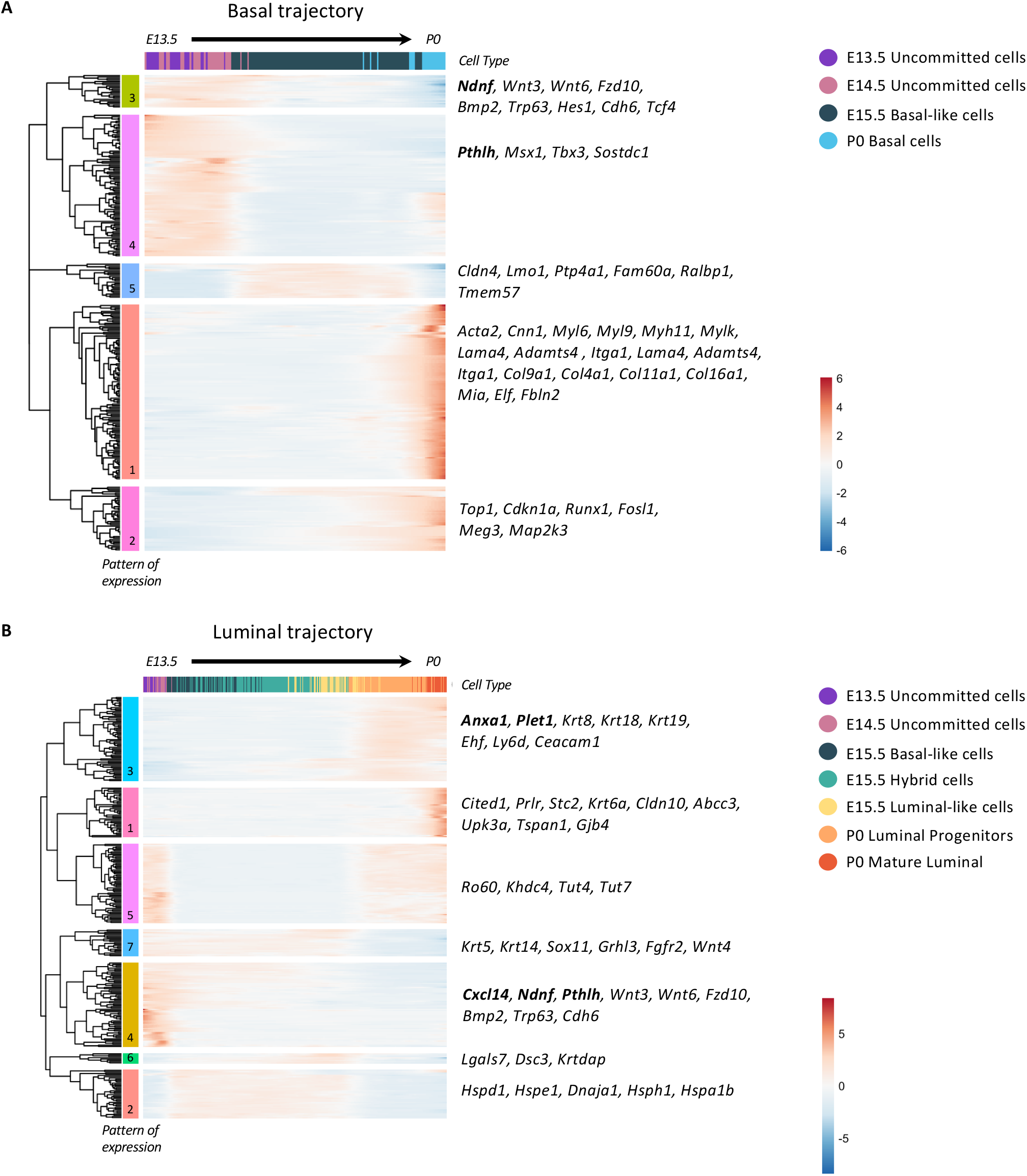
Pseudotime ordering identifies genes associated with early luminal and basal differentiation. (A and B) Heatmaps illustrating genes exhibiting a differential pattern of expression along the pseudotime (from E13.5 to P0) towards the basal lineage (A) or the luminal lineage (B). Genes (rows) are clustered based on the dendrogram on the left and color-coded by their expression levels (from blue to red). The gene expression levels were smoothed using the GAM and scaled by row. Genes of interest are indicated on the right. Each set of genes with a specific pattern is color-coded on the left: 5 distinct patterns in the basal lineage (A) and 7 unique patterns in the luminal lineage (B).

On the basal trajectory we found 5 distinct patterns of expression. Patterns 3 and 4 contained genes with sustained increased expression in early embryonic developmental times, at E13.5 and E14.5. Known key regulators of mammary bud epithelial cells are highly expressed only during early embryonic development, including *Ndnf, Pthlh, Msx1, Tbx3, Sostdc1*, whose expression is lost before birth. Moreover, multiple Wnt related genes, such as *Wnt3, Wnt6* and *Fzd10*, were enriched at these early developmental stages.

A different subset of genes, mostly related to cell migration (*Ptp4a1, Fam60a, Ralbp1*), appeared to be transiently upregulated at E15.5 (Pattern 5). Transcripts involved in mammary basal differentiation were progressively increasing towards the P0 basal cluster (Pattern 1); these included myosin-related proteins (*Myl6, Myl9, Myh11, Mylk*) and genes associated to ECM composition and organization (*Lama4, Adamts4, Itga1, Col9a1, Col4a1, Col11a1, Col16a1*). In addition, towards the P0 basal cluster, we also found increased expression levels of genes regulating cell proliferation (*Top1, Cdkn1a, Runx1, Fosl1*), cytoskeletal organization (*Tuba1c, Tubb6*) and angiogenesis (*Tnfrsf12a, Serpine1, Tgfa, Hbegf*) in Pattern 2, suggesting that epithelial growth is highly regulated at this developmental stage.

On the other hand, we observed 7 distinct expression patterns along the luminal differentiation trajectory. As expected, the pattern exhibiting increasing expression across the mammary developmental trajectory contains genes with known luminal characteristics, such as *Krt8, Krt18* and *Krt19* (Pattern 3). A second group of genes that is switched on during late stages of differentiation is enriched for ML cells markers, such as *Cited1* and *Prlr* (Pattern 1). Genes expressed at the beginning of the differentiation process and subsequently repressed along the luminal trajectory include typical basal markers, such as *Krt5* and *Krt14* (Pattern 7). *Sox11* also presents this dynamic pattern of expression, gradually decreasing along the differentiation process. Indeed, *Sox11* is expressed in MECs only during the early stages of MG embryonic development – when MG epithelial cells are largely quiescent – and is no longer detected by E16.5, consistent with our results. Of interest, *Sox11* has been recently involved in cell fate regulation in the embryonic MG (Tsang et al., 2021). Genes involved in epithelial stratification, such as *Lgals7, Dsc3* and *Krtdap*, are switched on only in luminal-like cells present at E15.5 (Pattern 6). Finally, Pattern 2 comprises genes encoding for several Heat shock proteins (Hsps). There is growing evidence that Hsps may impact neurodevelopment through specific pathways regulating cell differentiation, migration or angiogenesis (Miller & Fort, 2018).

To investigate whether lineage bias is reflected by spatial segregation of cells acquiring luminal or basal characteristics during embryonic development, we first identified genes that exhibited a lineage-specific expression pattern along the differentiation trajectories (Figure 3A-C). These included *Cxcl14, Ndnf* and *Pthlh* (Figure 2A) and *Anxa1, Plet1* and *Lgals3* (Figure 2B and S2E) for basal and luminal lineage specification respectively. Using single molecule RNA-fluorescence *in situ* hybridization (smRNA-FISH), we subsequently examined the spatiotemporal expression pattern of selected genes at distinct stages of mammary embryonic development. Probes for the luminal specific membrane-associated protein Annexin A1 (*Anxa1*) (Fankhaenel et al., 2021) and the basal-specific secreted chemokine *Cxcl14* (Sjöberg et al., 2016) revealed that at early embryonic stages (E13.5), *Cxcl14* is expressed in all MECs, and *Anxa1* is lowly expressed in rare cells homogeneously distributed within the mammary bud (Figure 3D). However, at the critical developmental time of E15.5, the transcripts for these two genes show divergent spatial distribution patterns, with *Anxa1* expression being mainly confined to cells in the inner bud region and *Cxcl14* transcripts restricted to the external cell layers in contact or close proximity with the BM (Figure 3D). By P0, *Anxa1* and *Cxcl14* showed clear luminal and basal restricted expression respectively (Figure 3E). To quantify the spatial segregation of gene expression, we divided the mammary bud into three concentric “rings” (outer, middle and internal regions) (Figure S2A) and counted the number of RNA molecules (represented by each dot) within each ring for both markers. This unbiased approach confirmed the uniform expression pattern of *Anxa1* and *Cxcl14* transcripts in all 3 regions of the mammary bud at E13.5 (Figure 3F). By E15.5, however, *Anxa1* transcripts were prominently restricted to the middle and inner ring, while *Cxcl14* transcripts appeared preferentially localized to the middle and outer ring of the mammary bud (Figure 3F). This was particularly intriguing as all MECs still express K5 (in white in Figure 3D, S2B-C) and other known markers of adult LCs and BCs at this developmental stage (Figure S2D). Thus, *Anxa1* and *Cxcl14* represent novel markers of MECs committed to luminal and basal lineages, respectively, as early as E15.5 during mammary development. Analogous smRNA-FISH analysis of E15.5 mammary buds with additional probes suggested that *Ndnf* and *Pthlh* are also expressed in embryonic basal committed MECs, while *Plet1* and *Lgals3* expression likely mark cells biased towards the luminal lineage (Figure S2B-C), further corroborating our temporal scRNA-seq analysis (Figure 2A-B).

**Figure 3.**
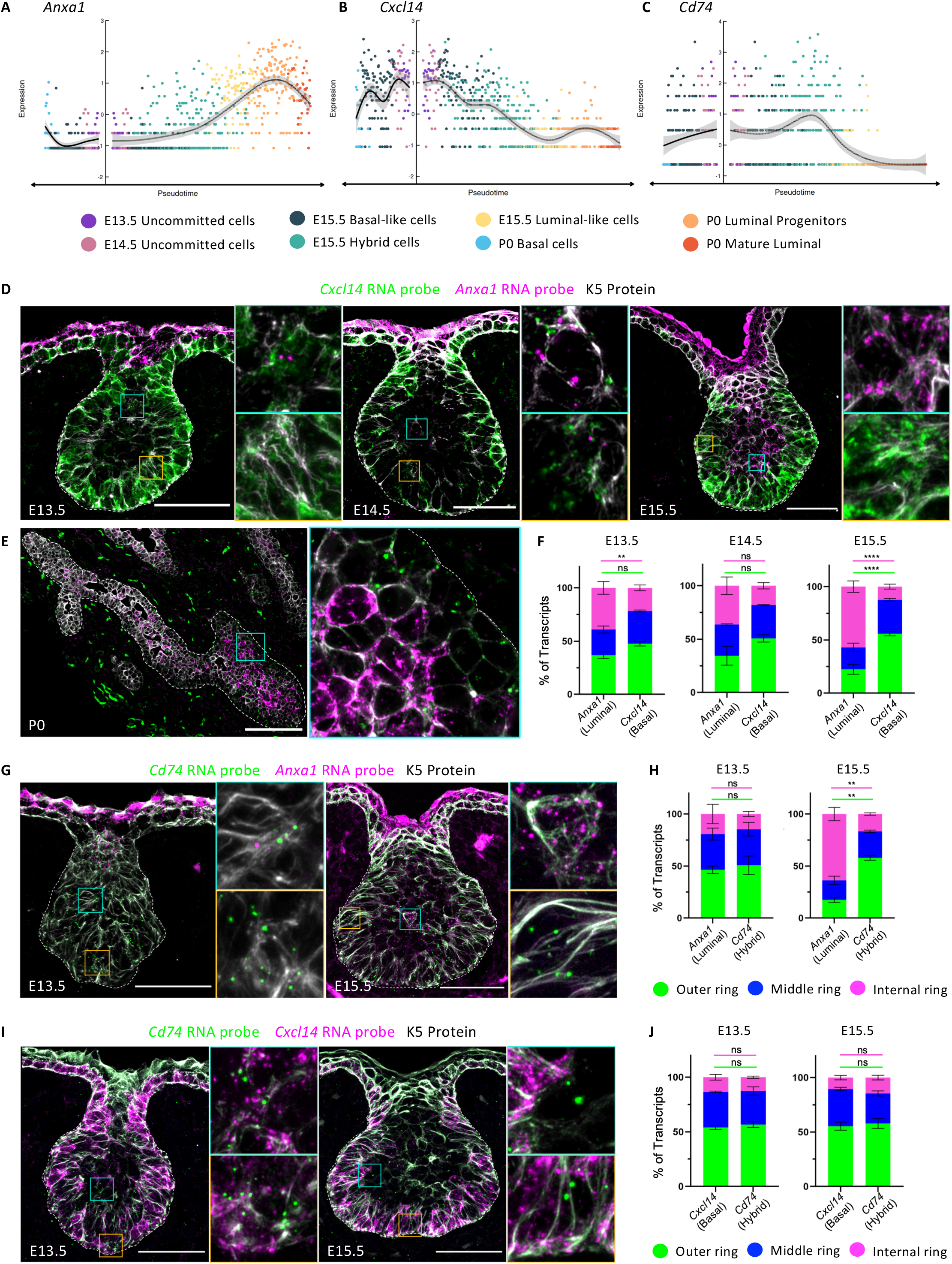
Luminal and basal progenitors are already physically separated at E15.5. (A-C) Examples of genes with pseudotime-dependent expression towards luminal differentiation (*Anxa1*, A), basal differentiation (*Cxcl14*, B) or with a higher expression in the hybrid cluster at E15.5 (*Cd74*, C). Cells are color-coded by cluster. (D and E) Representative sections of embryonic mammary buds at E13.5, E14.5 and E15.5 (D) and P0 (E) showing the expression of *Cxcl14* (in green) and *Anxa1* (in magenta) detected by RNAscope and immunostained with K5 (in white). Dotted lines delineate the BM. Scale bars: 50 μm (D), 100 μm (E). (F) Quantification of the proportion of *Anxa1* and *Cxcl14* transcripts in each ring at each developmental stage. (G) Representative sections of embryonic mammary buds at E13.5 and E15.5, showing the expression of *Cd74* (in green) and *Anxa1* (in magenta) detected by RNAscope and immunostained with K5 (in white). Dotted lines delineate the BM. Scale bar: 50 μm. (H) Quantification of the proportion of *Cd74* and *Anxa1* transcripts in each ring at each developmental stage. (I) Representative sections of embryonic mammary buds at E13.5 and E15.5 showing the expression of *Cd74* (in green) and *Cxcl14* (in magenta) detected by RNAscope and immunostained with K5 (in white). Dotted lines delineate the BM. Scale bar: 50 μm. (J) Quantification of the proportion of *Cd74* and *Cxcl14* transcripts in each ring at each developmental stage. Statistical significance in (F), (H) and (J) was assessed with two-way ANOVA test between the two probes. The statistical analysis was performed between the outside ring (green line) and the inside ring (magenta line). ns: non-significant, ** indicates p<0.01 and **** indicates p<0.0001.

In light of our findings that a proportion of MECs are already lineage committed at E15.5, we next sought to examine the spatial localization of cells possessing a hybrid basal-luminal expression signature within the developing mammary bud. To this aim, we searched for genes associated with the hybrid cell cluster identified at E15.5 (Figure 1B). A promising candidate marker gene for this cluster was the HLA class II cell surface receptor *Cd74* (Figure 3C, S2E), previously proposed as a putative mammary stem cell marker (dos Santos et al., 2013). smRNA-FISH analysis revealed that, while *Cd74* expression overlapped with both *Anxa1* and *Cxcl14* in early mammary embryonic development (E13.5), the vast majority of *Cd74* transcripts resided in the middle and outer regions of the mammary bud at E15.5, coinciding with *Cxcl14* expression (Figure 3G-J). Thus, the hybrid cells identified by transcriptomic analysis at E15.5 appear to be primarily localized in proximity with the BM, where basal-committed cells are also found within growing mammary buds.

Collectively, our spatial transcriptomic data reveal that the embryonic basal-like and luminal-like mammary cell clusters identified by scRNA-seq are already located in defined and mutually exclusive positions within the mammary bud at E15.5, at the onset of branching morphogenesis. Spatial segregation of mammary embryonic progenitors may conceivably underlie their state of differentiation and lineage commitment at this critical stage of embryonic mammary development.

### Identification of two spatially distinct mesenchymal cell populations in the embryonic mammary stroma

Mammary epithelial buds at E13.5 are surrounded by a specialized mammary mesenchyme, subsequently undergoing sprouting to invade the underlying fat pat precursor at around E15.5 to initiate the first stages of branching morphogenesis. Paracrine signaling between mammary epithelial and surrounding mesenchymal cells is indispensable for this process (Spina & Cowin, 2021; Wansbury et al., 2011). To gain further insights into mammary mesenchymal patterning during embryonic development, we focused our analysis on the scRNA-seq data of mesenchymal cells at E13.5, E15.5 and P0. Clustering of non-epithelial cells identified three mammary mesenchymal cell subsets at each stage (Figure 4A). By computing a cell cycle score based on a list of cell cycle-related genes, we identified proliferative cell clusters exclusively at early developmental timepoints, E13.5 and E15.5 (Figure S3A), indicating that proliferative cell populations are mostly absent at birth.

**Figure 4.**
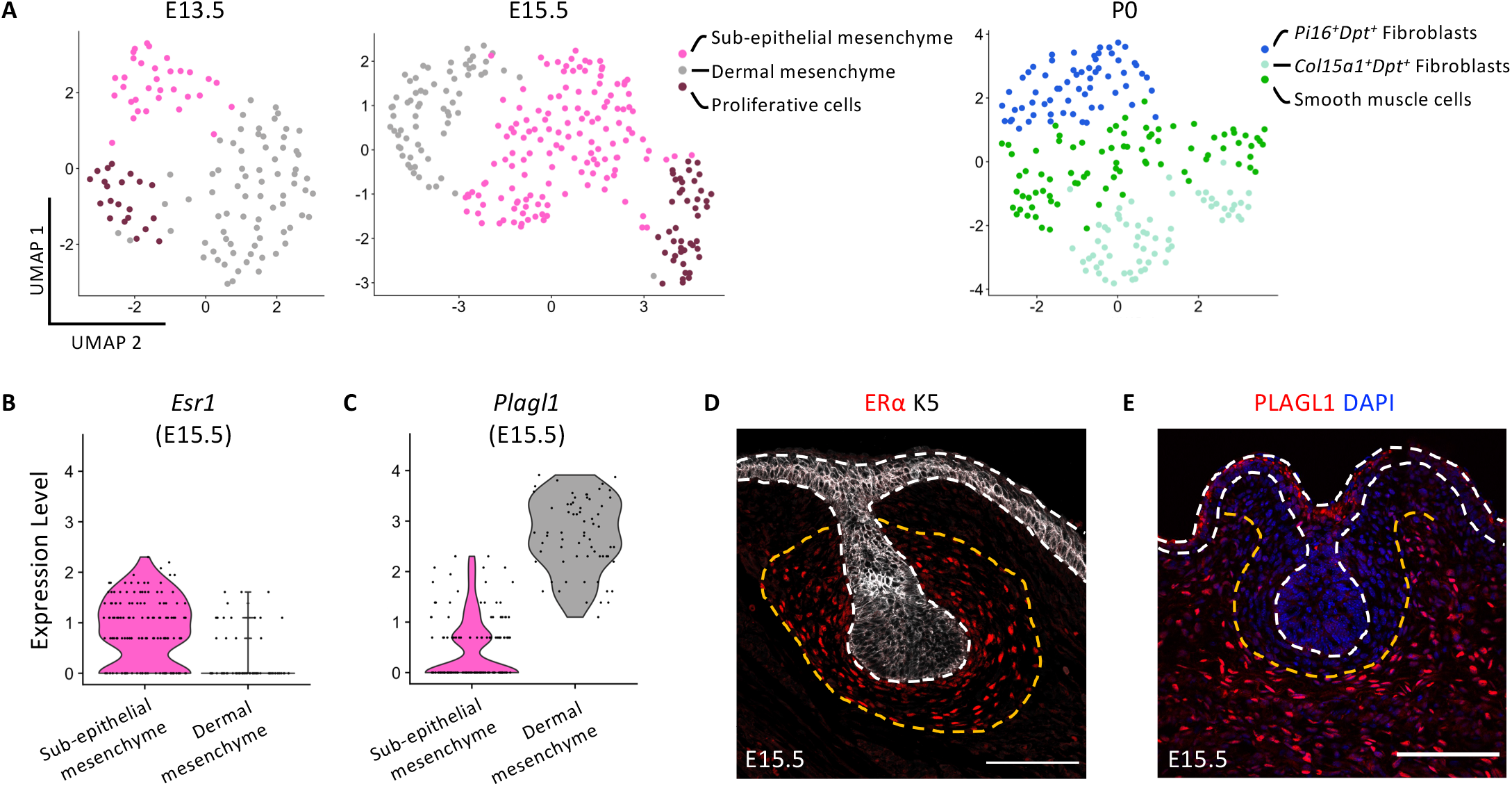
The embryonic mammary mesenchyme contains two spatially distinct cell populations. (A) UMAP plots of embryonic mammary mesenchymal cells isolated at E13.5, E15.5 and P0 after subset analysis. Cells are color-coded by cluster. (B and C) Violin plots representing the expression levels of *Esr1* (B) and *Plagl1* (C) in sub-epithelial and dermal mesenchyme respectively, at E15.5. (D and E) Representative sections of embryonic mammary buds at E15.5 immunostained for ERα (in red) and K5 (in white) (D) or PLAGL1 (in red) and DAPI (in blue) (E). Dotted lines delineate the BM (in white) and the two mesenchymal compartments (in orange). Scale bars: 100 μm.

We next singled out specific markers defining the two non-proliferative mesenchymal clusters at E15.5 (Figure S3B). Candidate genes included *Esr1* (coding for the ERα) and *Plagl1* (coding for the zinc finger protein PLAGL1), which were highly expressed in opposing mesenchymal clusters (Figure 4B-C, S3B). Immunostaining for ERα showed clear expression in mesenchymal cells directly surrounding the mammary bud (Figure 4D), as previously reported (Wansbury et al., 2011). Immunofluorescence analysis for PLAGL1, on the other hand, revealed that PLAGL1^+^ mesenchymal cells are located further away from the mammary epithelium (Figure 4E). These results suggest that the two transcriptionally distinct mesenchymal populations are also differentially localized within the embryonic mammary stroma, and can be categorized based on their proximity to the mammary epithelial bud. We thus refer to cells closest to the epithelium as the sub-epithelial mesenchyme and those located further away as dermal mesenchyme.

The heterogeneity of mesenchymal cells and the complexity of the mammary stroma increases at birth, where two clusters of *Dpt*^*+*^ fibroblasts can be distinguished, namely *Col15a1*^*+*^ and *Pi16*^*+*^ clusters, as previously identified across 17 other tissues (Buechler et al., 2021). Interestingly, the *Col15a1*^*+*^*Dpt*^*+*^ population also expresses *Fabp4, Pparg* and *Aoc3*, surface markers of pre-adipocytes. Conversely, the *Pi16*^*+*^*Dpt*^*+*^ population expresses *Dpp4, Sema3c* and *Wnt2*, which are reported to be upregulated in subcutaneous mesenchymal progenitors (Merrick et al., 2019) (Figure 4A, S3C). Structural and matricellular proteins of the ECM (*Col4a1, Col4a2, Col18a1, Mmp19, Sdc1, Sparcl1*) are also highly expressed in the *Col15a1*^*+*^*Dpt*^*+*^ population. Finally, the third mesenchymal population identified at P0 displays elevated expression of *Eln, Mfap4, Mgp*, genes typically expressed by smooth muscle cells.

### FGF10 produced by the dermal mesenchyme is an important regulator of embryonic mammary morphogenesis

Communication between the mammary epithelial and stromal compartment is essential for branching morphogenesis (Inman et al., 2015). Thus, in light of the observed spatial patterning of mesenchymal cells at E15.5 (Figure 4), we next sought to computationally predict specific paracrine interactions between the identified mesenchymal cell subsets and MECs using CellPhoneDB, a bioinformatic tool designed to predict highly significant ligand-receptor interactions between two cell types from scRNA-seq data (Vento-Tormo et al., 2018). We focused on ligand-receptor interaction pairs between the sub-epithelial or dermal mesenchyme and the basal-like cluster of MECs at E15.5, which we established to be in direct contact or in close proximity to the BM (Figure 3D-E). This approach highlighted several developmental signaling pathway components, including FGF, Wnt and Notch receptors and ligands, as putative mediators of the cross-talk between E15.5 basal-like cells and the sub-epithelial or dermal mesenchyme (Figure S4). Of particular interest, specific interactions between the FGFR2 and its soluble ligand FGF10, as well as between the Transforming growth factor beta receptors TGFBR1 and TGFBR2 and their ligand TGFB2 were highly significant between basal-like MECs and the more distant dermal mesenchymal cells (Figure S4). To functionally assess the validity of this computational prediction, we sought to investigate the impact of exogenous FGF10 on embryonic branching morphogenesis by live cell imaging of mammary buds established in *ex vivo* cultures. Explant cultures provide a highly tractable system for modelling embryonic mammary cell behavior and branching morphogenesis (Carabaña & Lloyd-Lewis, 2022; Voutilainen et al., 2013). Embryonic mammary buds along with their surrounding mesenchyme were dissected at E13.5 and cultured *ex vivo* on an air-liquid interface. Embryonic MECs expressed both basal and luminal markers (K5, K14 and P63 for basal cells and K8 for luminal cells) after 24h in culture (Figure S5A-C), consistent with *in vivo* observations (Figure S2D) (Wansbury et al., 2011). During 8 days of *ex vivo* culture, embryonic mammary buds undergo sprouting and branching, recapitulating the morphogenetic events occurring *in vivo* (Figure S5D-E). Immunostaining of the resulting 8-day-old ductal tree (corresponding to approximately P0/P1 *in vivo*) revealed that MECs in the outer layer express basal markers such as P63 (Figure S5D, S5F) and smooth muscle actin (α-SMA) (Figure S5E), while inner layer cells express the luminal marker K8 (Figure S5D-E). In addition, polarity acquisition appeared normal, as revealed by apical ZO-1 staining in the inner layer of luminal cells (Figure S5F). Thus, key aspects of embryonic mammary morphogenesis and epithelial lineage segregation can be reconstituted in *ex vivo* cultures.

Taking advantage of this powerful system, we next investigated the impact of FGF signaling by undertaking live-imaging of embryonic mammary explants cultured with FGF10 (Figure 5A). To measure the velocity of branch growth in control and FGF10 treated conditions, after 4 days in culture we traced the endpoint of each branch acquired every 60 min for 24 hr. By measuring the distance travelled over time in control and FGF10 treated conditions, these experiments indicated that mammary branches grow faster when cultured in the presence of FGF10 (Figure 5B).

**Figure 5.**
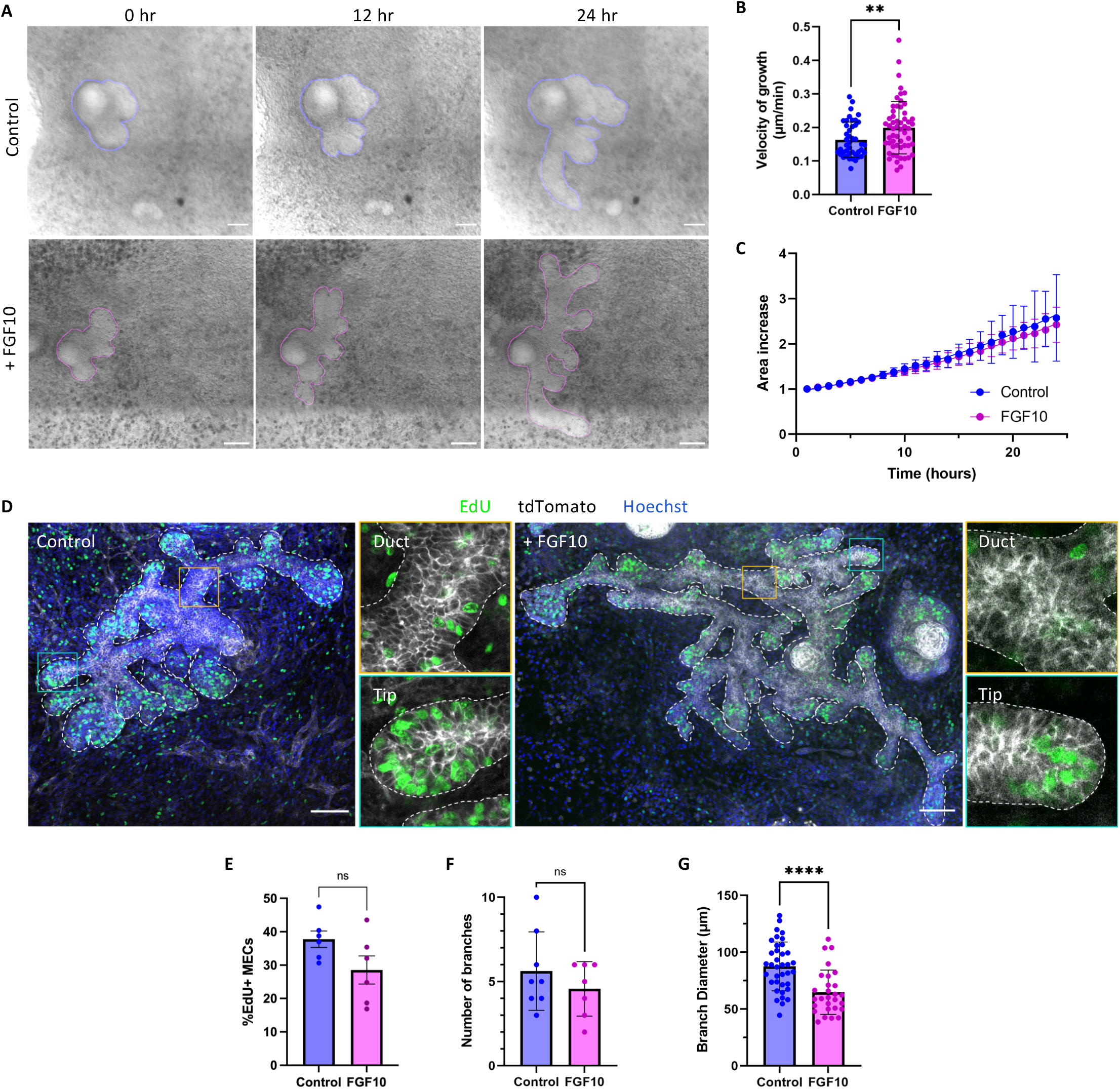
FGF10 accelerates embryonic mammary branching without affecting cell proliferation. (A) Time-lapse images of a mammary explant grown in control medium (top) or in the presence of FGF10 (bottom) for 24 hr. T= 0h refers to 4 days in culture. Scale bars: 100 μm. The rendered surface of the mammary epithelium is outlined in blue (in the control bud) and in magenta (in the FGF10 condition). (B) Quantification of the velocity of branch growth in control conditions (n= 43) and in the presence of FGF10 in the medium (n= 56). (C) Fold change increase in area in control and FGF10 conditions. In both cases, the area is doubled within 16 hr in culture. (D) Representative whole-mount immunostaining of an embryonic mammary gland cultured in control and FGF10 conditions showing Edu^+^ cells (in green), membrane tdTomato (in white) and DAPI (in blue). Mammary buds were dissected at day E13.5 and cultured *ex vivo* for 7 days. Orange outlined insets show a duct region and blue outlined insets show a tip region. (E, F and G) Quantification of Edu^+^ cells (E), number of branches (F) and branch diameter (G) in control and FGF10 conditions. Statistical significance was assessed with two-tailed unpaired Welch’s t-test. ** p< 0.01, **** p<0.0001, ns: non-significant.

Mesenchymal-produced FGF10 may accelerate branching morphogenesis by increasing either epithelial cell proliferation or motility. To discriminate between these two possibilities, we measured the planar surface area of mammary buds over time and found that tissue growth was not significantly affected by FGF10, since the explant area increased 2-fold within 16 hours of culture in both control and FGF10 conditions (Figure 5C). While FGF10 is a potent mitogen in several contexts, 5-ethynyl-2′-deoxyuridine (EdU) incorporation experiments suggested that it did not promote mammary epithelial cell proliferation during branch elongation in *ex vivo* cultures (Figure 5D-E). Moreover, the number of branches in embryonic explant cultures supplemented with FGF10 was equivalent to control cultures (Figure 5F). However, the diameter of branches at their base was reduced in the presence of FGF10 (Figure 5G), suggesting that while MEC numbers are equivalent, cells may move faster along extending ducts, which consequently become thinner in the presence of FGF10. Our data therefore shows that, similar to observations made during pubertal branching morphogenesis (Hannezo et al., 2017), FGF signaling promotes branching of the embryonic mammary ductal tree at the initial stages of embryonic development, likely by promoting epithelial cell motility.

## Discussion

To generate complex organs of diverse shapes and function, tissue morphogenesis and cell fate specification must be tightly coordinated. Yet, how morphological changes steer individual cells towards a particular fate and, conversely, how cell fate decisions orchestrate morphogenesis, remain ambiguous. By combining temporal scRNA-seq analysis with spatial transcriptomics and live imaging of tissue explant cultures, this work provides new insights into the progressive lineage specification of epithelial cells during embryonic mammary ductal development.

Our data revealed that embryonic MECs at E15.5 can already be distinguished as three transcriptionally discrete cell populations: basal-like cells, luminal-like cells and ‘hybrid’ cells. This was surprising, as previous scRNAseq studies concluded that bipotent MaSCs, sharing luminal and basal characteristics, exist throughout embryogenesis, and the two separate lineages are only specified postnatally (Giraddi et al., 2018; Wuidart et al., 2018). The high quality and depth of sequencing attained in this study, however, likely enabled us to identify different embryonic MEC clusters that were previously indistinguishable by gene expression. Indeed, more recent snATAC-seq analysis of E18.5 and adult MG revealed that E18.5 MECs, although still presenting fetal-specific features, are partially lineage-biased and already harbor adult-like basal, LP and ML characteristics (Chung et al., 2019). The results presented herein are also consistent with our previous lineage tracing and theoretical modeling analyses (Lilja et al., 2018), which implied that lineage potential restriction coincides with the initiation of branching morphogenesis around E15.5. Collectively, our data supports a model whereby these two processes are linked. As cells rearrange their position within the growing tissue, coordination between cell differentiation and cell movements may be mediated by their exposure to changing environmental cues. By determining the regional positioning of the different cell clusters that we identified by scRNAseq, we observed that luminal and basal commitment is indeed reflected by differences in cell localization within the developing mammary epithelium. It is conceivable, therefore, that spatial segregation of mammary embryonic progenitors at this critical stage of development underpins their state of differentiation and lineage commitment.

Based on these results, we propose a dynamic hierarchical model of mammary differentiation spanning embryonic development (Figure 6). Mammary epithelial cells at E13.5 are undifferentiated and have yet to engage in lineage specification. As development and tissue morphogenesis progress, these putative multipotent embryonic MaSCs will first give rise to basal-like cells, designated as such based on their expression of several genes that define basal mammary cells postnatally. Basal-like cells will then either differentiate into basal unipotent progenitors by P0, or they will transition towards a transcriptionally hybrid state. Hybrid cells, whose lineage potential remain unclear at this stage, will gradually lose basal markers concomitant with acquiring luminal gene expression, eventually giving rise to unipotent luminal cells at birth.

**Figure 6.**
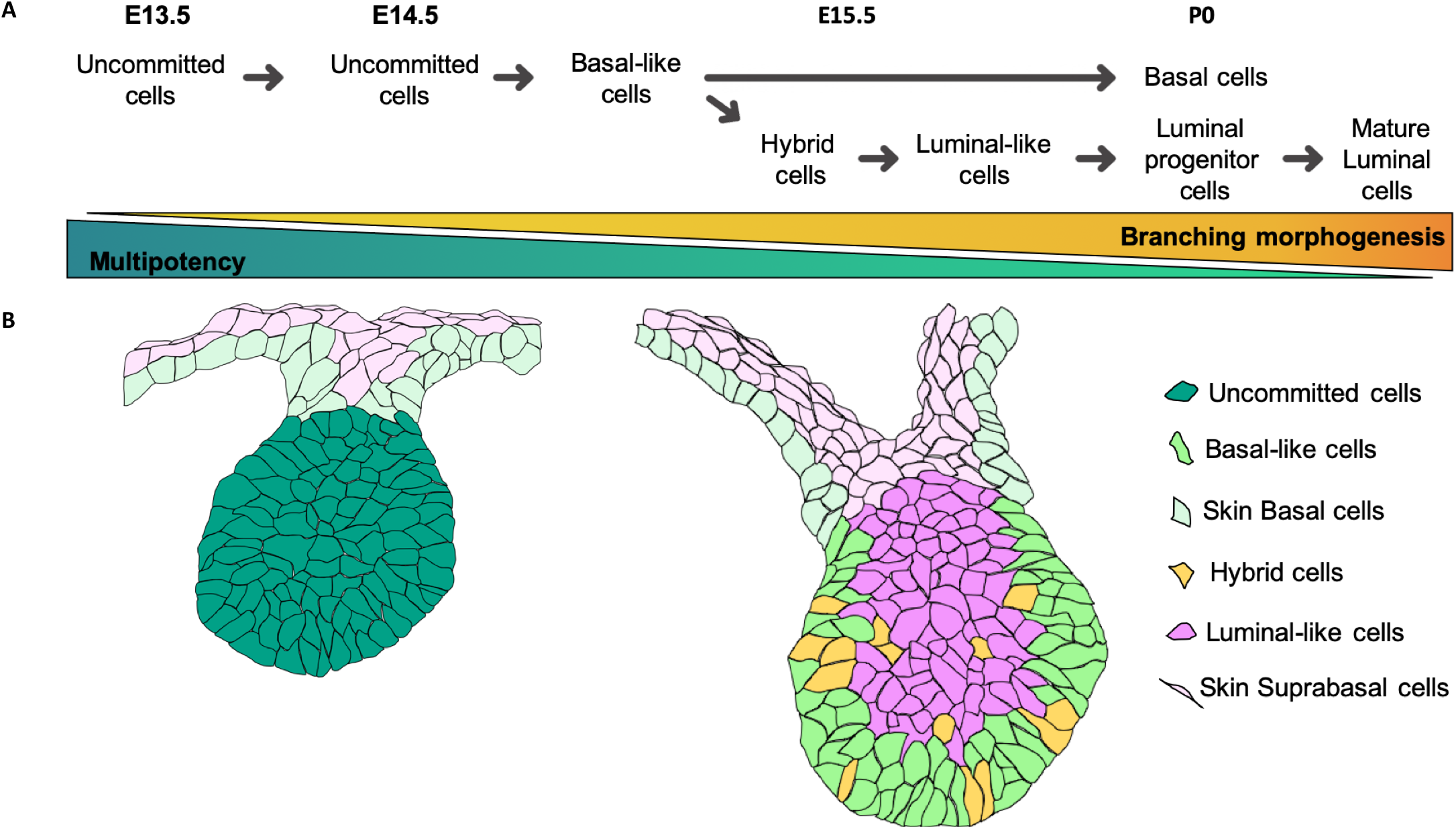
Proposed model for lineage segregation of embryonic mammary epithelial cells during development. (A) Proposed model of luminal and basal differentiation trajectories from E13.5 to P0. (B) Cartoon depicting the spatial localization of the different cell types distinguishable in the embryonic mammary bud at E13.5 and E15.5.

Embryonic MECs co-express the differentiation markers commonly used to distinguish LCs and BCs in the adult mammary gland (Figure S2D). This has, to date, hampered studies of the precise timing and molecular regulators of embryonic mammary lineage specification. The comprehensive single cell transcriptomic atlas compiled in this work enabled the spatial mapping of distinct subsets of embryonic mammary cells, some of which are already committed to basal or luminal fate. In addition to facilitating the *in situ* identification of potentially multipotent and unipotent mammary progenitors, the lineage-specific genes we discovered may be functionally important for dictating cell fate choices. These novel early markers of luminal or basal commitment likewise provide new specific promoters that could be used in future lineage tracing studies to definitively establish the differentiation dynamics and lineage potential of early mammary progenitors.

Additionally, our study provides important insights into the poorly explored resident mammary embryonic mesenchymal cell populations that direct epithelial branching morphogenesis. We identified specific transcriptional signatures that distinguish two spatially-restricted mesenchymal populations in mammary embryonic glands, named sub-epithelial and dermal mesenchyme. It remains unclear however how mesenchymal cells adopt a fibroblast or an adipocyte fate during embryonic development. Addressing this important question awaits future fate-mapping studies using specific stromal Cre drivers based on the promoters of genes identified in this work.

Ligand-receptor pair interaction analysis of the compiled scRNAseq data implicated several components of the FGF pathway as important mediators of communication between dermal mesenchyme and basal-like cells. Detailed scrutiny of differential gene expression in the sequenced mesenchymal embryonic cells, also revealed that the dermal mesenchyme contained cells expressing genes implicated in cell invasive behavior (*Cxcl12*) and axon guidance (*Nrp2, Epha3, Epha7*). These findings imply that dermal mesenchymal cells might secrete signaling factors that promote epithelial branching morphogenesis and fat pad invasion. In fact, knock-out mice for the FGF receptor *Fgfr2b* or its ligand *Fgf10* fail to develop mammary placodes, suggesting that FGF10-FGFR2B signaling is required to initiate embryonic mammary development (Mailleux et al., 2002). However, this phenotype precluded studies into the role of the FGF10-FGFR2B signaling axis on mammary embryonic development. E*x vivo* mammary embryonic explant cultures, by contrast, provides opportunities to overcome challenges associated with genetic knock-out models. Our live-imaging data and custom-made image analysis pipeline revealed that, in the presence of exogenous FGF10, embryonic mammary branching is accelerated. Whether this is associated with more rapid differentiation of mammary progenitors, however, warrants further investigation in future studies.

In summary, this work reveals the cell-state heterogeneity of the embryonic mammary epithelium and surrounding mesenchyme, and provides important insights into the paracrine interactions that guide branching morphogenesis. Our computational analyses have uncovered the molecular mechanisms and transcription factors involved in regulating mammary cell fate specification. Furthermore, the lineage trajectory analysis reported herein could be extended to other stratified epithelia to determine whether these mechanisms are shared in other organs during embryonic development.

## STAR Methods

### Mouse models

*Ex vivo* cultures were established from the double fluorescent reporter R26^mT/mG^ mice (Muzumdar et al., 2007) in a mixed genetic background. We exclusively analyzed female mice. WT C57B6 mice were analyzed at embryonic stages E13.5, E14.5 and E15.5, and during postnatal development at P0, as indicated in the figure legends. Plug detection at mid-day was considered 0.5 days-post-coitus (E0.5). Mice were genotyped by PCR analysis on genomic DNA extracted from an ear piece for adult mice or tail tip for embryos.

### Ethics Statement

All studies and procedures involving animals were in accordance with the recommendations of the European Community (2010/63/UE) for the Protection of Vertebrate Animals used for Experimental and other Scientific Purposes. Approval was provided by the ethics committee of the French Ministry of Research (reference APAFIS #34364-202112151422480). We comply with internationally established principles of replacement, reduction, and refinement in accordance with the Guide for the Care and Use of Laboratory Animals (NRC 2011). Husbandry, supply of animals, as well as maintenance and care in the Animal Facility of Institut Curie (facility license #C75–05–18) before and during experiments fully satisfied the animal’s needs and welfare. All mice were housed and bred in a specific-pathogen-free (SPF) barrier facility with a 12:12 hr light-dark cycle and food and water available *ad libitum*. Mice were sacrificed by cervical dislocation as adults or decapitated as embryos.

### Embryonic mammary gland dissection and *ex vivo* culture

Mammary embryonic buds were dissected following the protocol developed by the laboratory of M. Mikkola (Voutilainen et al., 2013). Briefly, embryos were harvested from the uterus of a pregnant dam at day E13.5 of pregnancy. Under a dissecting microscope, an incision along the dorsal-lateral line from the hind limb to the forelimb in the right flank of the embryo was done using spring scissors. The flank of the embryo from the incision along the dorsal-lateral line to the midline was detached and the same steps were repeated for the left flank of the embryo, but this time cutting along the dorsal-lateral line from the forelimb to the hind limb. Tissues were collected in a 24-well plate with phosphate buffered saline (PBS) until all embryos were dissected.

Next, proteolytic digestion of dissected embryonic flanks was performed as previously described (Lan & Mikkola, 2020). Tissues were incubated with freshly prepared 1.25 U/ml Dispase II solution (Roche, 04942078001) at 4°C for 15 minutes. Then, with Pancreatin-Trypsin solution at room temperature (RT) for 4-5 minutes. To prepare Pancreatin-Trypsin working solution: first 0.225 g of Trypsin (Sigma-Aldrich, 85450C) were dissolved into 9 mL of Thyrode’s solution [8 g/L NaCl (Sigma-Aldrich, S5886) + 0.2 g/L KCl (Sigma-Aldrich, P5405) + 0.05 g/L NaH_2_PO_4_ • H_2_O (Sigma-Aldrich, S3522) + 1 g/L D-(+)-Glucose (Sigma-Aldrich, G7021) + 1 g/L NaHCO_3_ (Sigma-Aldrich, S5761) dissolved in 1 L of distilled water and filter sterilized]. Then, 1 mL of 10X Pancreatin stock solution [0.85 g NaCl (Sigma-Aldrich, S5886) and 2.5 g Pancreatin (Sigma-Aldrich, P3292) dissolved into 100 mL of distilled water on a magnetic stirrer on ice for 4 hr and filter sterilized] and 20 μL of Penicillin-Streptomycin (10,000 U/ml in stock) (Sigma-Aldrich, P4333) were added. Finally, pH was adjusted to 7.4 with NaOH and the solution was filter sterilized (see in (Lan & Mikkola, 2020)).

When skin epithelium started to detach from the edges of the mammary mesenchyme, the Pancreatin-Trypsin solution was replaced with DMEM/F-12 (Gibco-Thermo Fisher Scientific, 21331020) embryonic culture medium to inactivate the enzyme activity. After incubating the tissue for 20-30 minutes in ice, the skin epidermis was removed away from the mesenchyme containing the embryonic mammary buds using two needles.

Mammary embryonic buds were established in *ex vivo* culture as previously detailed in (Carabaña & Lloyd-Lewis, 2022). Collected embryonic mammary tissue was placed on a cell culture insert floating on embryonic culture medium into a 35 mm cover glass-bottomed tissue culture dish (Fluorodish, 81158). Embryonic culture medium is DMEM/F-12 (Gibco-Thermo Fisher Scientific, 21331020) supplemented with 2 mM GlutaMAX™ (Gibco-Thermo Fisher Scientific, 35050-038), 10% fetal bovine serum (FBS) (v/v), 20 U/ml Penicillin-Streptomycin (Gibco-Thermo Fisher Scientific, 15140122) and 75 μg/mL Ascorbic acid (Sigma, A4544). Mammary cultures were maintained in a tissue culture incubator at 37°C with 5% CO_2_. The culture media was replaced every two days. For growth factor assays, 1 nM FGF10 (Bio-techne, 6224-FG) was added to the medium at day 4.

### Mammary cultures wholemount immunostaining

*Ex vivo* cultures whole-mount immunostaining was performed as previously described (Carabaña & Lloyd-Lewis, 2022). Explants were transfer to a 24 well plate, washed in PBS and fixed with 4% PFA for 2 hr at RT. After a blocking step in PBS containing 5% FBS, 1% Bovine Serum Albumin (BSA) and 1% Triton x-100 (Euromedex, 2000-C) for 2 hr, explants were incubated with primary antibodies diluted in blocking buffer overnight at 4°C. Then, with secondary Alexa-fluor conjugated antibodies and DAPI (10μM) diluted in PBS for 5 hr at RT. *Ex vivo* cultures were mounted in a slide using Aqua-Polymount (Polysciences, 18606). The following primary antibodies were used: rabbit anti-SMA (1:300, Abcam, ab5694), rat anti-K8 (1:300, Developmental Studies Hybridoma Bank, clone TROMA-I), mouse anti-P63 (1:300, Abcam, ab735), rabbit anti-K5 (1:300, Covance, PRB-160P-100), rat anti-ZO-1 (1:100, Millipore, MABT11), rabbit anti-K14 (1:300, Abcam, ab181595). Complete detail of the antibodies used here are provided in Key Resources Table 2.

EdU incorporation was visualized using Click-It chemistry (Invitrogen) by incubating *ex vivo* cultures for 2 hr with EdU solution (10 μM). EdU was then detected with freshly made Click-iT EdU Alexa Fluor 647 Imaging Kit (Invitrogen-Thermo Fisher Scientific, C10640), according to the manufacturer’s protocol. Nuclei were stained with Hoechst33342 (10 μg/mL) for 30 minutes at RT.

### Immunostaining on 2D sections

Embryos were harvested and fixed in 4% PFA overnight at 4°C, followed by another overnight incubation at 4°C in 30% sucrose. Then, embryos were embedded in optimum cutting temperature (OCT) compound and 7 μm-thick cryosections were cut using a cryostat (Leica CM1950). After a blocking step in PBS containing 5% FBS, 2% BSA and 0.2% Triton x-100 for 2 hr, sections were incubated with primary antibodies diluted in blocking buffer overnight at 4°C in a humidified chamber, then with secondary Alexa-fluor conjugated antibodies and DAPI (10μM) diluted in PBS for 2 hr at RT. Finally, sections were mounted in a slide using Aqua-Polymount (Polysciences, 18606). The following primary antibodies were used: rat anti-K8 (1:300, Developmental Studies Hybridoma Bank, clone TROMA-I), mouse anti-P63 (1:300, Abcam, ab735), mouse anti-ERalpha (1:20, Agilent-Dako, M7047), rabbit anti-K5 (1:300, Covance, PRB-160P-100), rabbit anti-PLAG1 (1:100) (Spengler et al., 1997). Complete detail of the antibodies used here are provided in Key Resources Table 2.

### Single molecule RNA fluorescence in situ hybridization (smRNA FISH)

smRNA-FISH was performed using the RNAscope Multiplex Fluorescent Reagent Kit v2 (Advanced Cell Diagnostics), according to the manufacturer’s recommendations. In brief, tissue cryosections were pre-treated with the target retrieval reagent (ACD, 322000) for 5 minutes and digested with Protease III (ACD, 322381) at 40°C for 15 minutes, before hybridization with the target oligonucleotide probes. Probe hybridization, amplification and binding of dye-labelled probes were performed sequentially. For subsequent immunostaining, sections were incubated in blocking buffer (PBS containing 5% FBS and 2% BSA) for 1 hr. For smRNA-FISH in *ex vivo* cultures, the blocking buffer also included 0,3% Triton x-100 (Euromedex, 2000-C) to allow tissue permeabilization. Incubation with primary antibodies diluted in blocking buffer was performed overnight at 4°C in a humidified chamber, then secondary antibodies and DAPI diluted in PBS were added for 2 hr at RT. The experiments were performed on at least three different embryos for each probe. Slides were mounted in ProLong Diamond Anti-fade Mountant (Invitrogen-Thermo Fisher Scientific, P36930) for imaging.

The following RNAscope probes were used: Mm-Anxa1-C2 (ACD, 509291), Mm-Lgals3-C2 (ACD, 461471), Mm-Plet1-C1 (ACD, 557941), Mm-Ly6d-C1 (ACD, 532071), Mm-Cxcl14-C3 (ACD, 459741), Mm-Ndnf-C2 (ACD, 447471), Mm-Pthlh-C3 (ACD, 456521), Mm-Cd74-C1 (ACD, 437501), 3-plex Positive Control Probe-Mm (ACD, PN 320881) and 3-plex Negative Control Probe (ACD, PN 320891). Complete details of RNAscope probes used here are provided in Key Resources Table 1.

### Microscopy and image acquisition

#### 3D imaging

Images were acquired using a LSM780 or LSM880 inverted laser scanning confocal microscope (Carl Zeiss) equipped with 25x/0,8 OIL LD LCI PL APO or 40x/1,3 OIL DICII PL APO. For standard 4-color imaging, laser power and gain were adjusted manually to give optimal fluorescence for each fluorophore with minimal photobleaching. Images were captured using the ZEN Imaging Software and processed in Fiji (ImageJ v1.53).

#### smRNA-FISH

images were acquired using a LSM880 with an Airyscan system. The Airyscan system has 32-channel GaAsP (Gallium Arsenide Phosphide) detectors, which allow to obtain images with enhanced spatial resolution and improved signal-to-noise ratio (SNR) than in traditional LSM systems (Huff, 2015). A 63x/1,4 OIL DICII PL APO objective was used. Images were processed in Fiji (ImageJ v1.53).

#### Live-imaging

time-lapse images were acquired using an LSM780 or LSM880 inverted laser scanning confocal microscope (Carl Zeiss) equipped with 10x/0,3 DICI EC PL NEOFLUAR, for imaging at the tissue scale. Explants were cultured in a humidified chamber at 37°C with 5% CO_2_ during the course of imaging. To analyze branching morphogenesis in embryonic mammary buds, images were acquired at 8 mm Z intervals over approximately 80 mm thickness and 60 min intervals for 12-48 hr.

### Single cell dissociation of embryonic mammary gland

The isolated embryonic mammary rudiments include both the mammary epithelium and the surrounding mesenchyme. 60-90 mammary rudiments were dissected for each experiment from 7-12 female embryos derived from 2-4 timed pregnant females. The scRNA-seq of each developmental time was performed in a separate dissection session to maximize the number of mammary buds analyzed/timepoint.

Embryonic mammary buds along with their surrounding mesenchyme were dissected as detailed above (see *Embryonic mammary gland dissection and ex vivo culture* section). Single cell dissociation was performed as previously described (Wuidart et al., 2018) with the following modifications:

- for mammary rudiments at E13.5, E14.5 and E15.5, single cell dissociation was performed through enzymatic digestion with 300 U/ml collagenase A (Roche, 10103586001) and 300 U/ml hyaluronidase (Sigma, H3884) for 90 minutes at 37°C under shaking. Mammary rudiments from each female embryo were dissociated in a separated 2 mL protein LoBind tube (Eppendorf, 022431102). Cells were further treated with 0.1 mg/ml DNase I (Sigma, D4527) for 3 minutes. 10% FBS diluted in PBS was added to quench the DNase I. Cells were pelleted by centrifugation at 320 g for 10 minutes.
- for mammary glands at birth, the enzymatic digestion for single cell dissociation was optimized as followed. 600 U/ml collagenase A (Roche, 10103586001) and 150 U/ml hyaluronidase (Sigma, H3884) for 90 minutes at 37°C under shaking were used for enzymatic digestion. Cells were further treated with 0.1 mg/ml DNase I (Sigma, D4527) for 3 minutes and an additional incubation in 0.63% NH_4_Cl for 1 minute allowed lysis of red blood cells. Cells were pelleted by centrifugation at 320 g for 10 minutes.

For all developmental times, after careful removal of the supernatant, cells were incubated in fluorescently labelled primary antibodies.

### Cell labelling, flow cytometry and sorting

Single cell suspensions were incubated for 15 minutes on ice with fluorescently labelled primary antibodies diluted in HBSS with 2% FBS. Cells were washed from unbound antibodies with 2% FBS in HBSS and the cell suspension was passed through a 40 μm cell strainer filter to eliminate cell clumps.

Cell viability was determined with DAPI and doublets were systematically excluded during analysis. CD45^+^, CD31^+^, Ter119^+^ (Lin^+^) non-epithelial cells were excluded. FACS analysis was performed using an ARIA flow cytometer (BD).

The following primary antibodies were used at a 1:100 dilution: APC anti-mouse CD31 (Biolegend, 102510), APC anti-mouse Ter119 (Biolegend, 116212), APC anti-mouse CD45 (Biolegend, 103112), APC/Cy7 anti-mouse CD49f (Biolegend, 313628), and PE anti-mouse EpCAM (Biolegend, 118206). The isotype controls were the following: PE rat IgM (Biolegend, 400808), PE/Cy7 rat IgG2a (Biolegend, 400522), APC/Cy7 rat IgG2a (Biolegend, 400524) and APC rat IgG2b (Biolegend, 400612). Complete details of the antibodies used are provided in the Key Resources Table 1. The results were analyzed using the FlowJo software (V10.0.7).

### Image analysis and quantification

For time-lapse live imaging analysis, first time-lapse reconstructions were generated using the Bio-Formats plugin (Linkert et al., 2010) in Fiji (ImageJ v1.53). Then, automated segmentation of mammary buds was performed using a custom-made segmentation model based on U-Net (Ronneberger et al., 2015). Segmented masks and raw image were input in the ImageJ plugin, BTrack, for tracking the growing branch tips. BTrack allows the users to remove or create new end points to manually correct the obtained tracks. We obtained the average growth rate for each branch using customized Python scripts (see Data and code availability). Statistical analyses were performed in Prism (v9.2, GraphPad).

To determine bud surface area in the presence of FGF10 in the medium, segmented masks were obtained from each timepoint using the U-Net model previously described. Generated masks were manually checked and corrected against raw data for consistency prior to extracting area measurements. Surface area was measure for each timepoint and statistical analyses were performed in Prism (v9.2, GraphPad).

For smRNA-FISH dot counting, the Find Maxima tool in Fiji (ImageJ v1.53) was used to find the highest peak values in the images using a previously specified threshold. Then, a custom ImageJ macro was coded to create 3 parallel regions of interest (ROIs) with a ring-shaped surface. Finally, the number of dots in each ROI was calculated for each smRNA-FISH probe. The percentage of dots in each ring was calculated as the ratio of number of dots in a specific ROI to the total number of dots in the 3 ROIs (outer, middle and internal ring). Statistical analysis was performed in Prism (v9.2, GraphPad).

For EdU quantification 3 independent explants in each condition were analyzed. For each explant, independent regions of interest were randomly selected in discrete Z-slides. The mammary epithelium was outlined manually in Fiji using the tdTomato or luminal lineage marker staining as a guide (ImageJ v1.53). Hoechst images were processed with a median filter (1-2px). StarDist (Schmidt et al., 2018; Weigert et al., 2020) was used to segment and quantify number of Nuclei and EdU^+^ nuclei within the outlined mammary epithelial tree region in Fiji (ImageJ v1.53). EdU^+^ nuclei were expressed as a percentage over total number of nuclei. Statistical analysis was performed in Prism (v9.2, GraphPad).

### scRNA-seq data processing and cluster analysis

Single cell capture and library construction were performed using the 10x Genomics Chromium Single Cell 3’ v3.1 kit following the manufacturer’s instructions, for samples of different developmental stages. The libraries were sequenced with an Illumina NovaSeq 6000 sequencer by the *Next Generation Sequencing* platform of Institut Curie.

#### Data pre-processing and quality control

The 10x Genomics Cell Ranger Single-Cell Software Suite was used for demultiplexing, read alignment and unique molecular identifier (UMI) quantification (http://software.10xgenomics.com/single-cell/overview/welcome). The pre-built mm10 reference genome obtained from the 10X Genomics website was used to align the reads. Then, the count matrices were individually loaded for each sample in R and analyzed using the Seurat package v4.0.5 (Hao et al., 2021).

Genes expressed in less than 3 cells and cells with UMI count < 5000 and mitochondrial UMI count > 6% were removed. This resulted in the following total number of high-quality cells: 228 at E13.5, 59 at E14.5, 740 at E15.5, 409 at P0 in WT mice.

#### Normalization

Objects were normalized separately using the SCTransform method, implemented in the “SCTransform’’ function from Seurat. Briefly, this method regresses out the sequencing depth variation between cells using a negative binomial regression model with regularized parameters (Hafemeister & Satija, 2019).

#### scRNA-seq data dimension reduction and clustering

Principal Component Analysis (PCA) was performed on the top 2000 highly variable genes of the SCT assay from the “SCTransform” step. The top 15 principal components (PCs) were further selected (based on inspection of PC elbow plot) to perform graph-based clustering and cell cluster detection. All the Uniform Manifold Approximation and Projection (UMAP) plots (McInnes et al., 2018) were computed using the “RunUMAP” Seurat function with default Seurat parameters.

#### Cell cluster identification

Cell clustering was performed using a two-step wise approach, using the “FindNeighbors’’ and “FindClusters’’ functions, respectively. The “FindClusters” function was used to set the resolution parameter to 0.8.

#### Differential expression analysis

Cell-type marker genes for each cluster were identified using the function “FindAllMarkers” function in Seurat, with detected in minimum cell fraction > 10% and log-fold change > 0.1. Then, cell clusters were manually annotated based on cell type specific markers that are known to be enriched in each cell population. Cell proliferative clusters were identified by using the following list of genes: ‘Pclaf’, ‘Ncapg2’, ‘Smc2’, ‘Tyms’, ‘Tuba1b’, ‘Hmgb2’, ‘Top2a’, ‘Tacc3’, ‘Cenph’, ‘Cdk1’, ‘Tubb5’, ‘Diaph3’, ‘Cenpf’, in order to compute an expression score using the Seurat function ‘AddModuleScore’.

#### Signature construction

a single-cell ID score for “basal-like” and “luminal-like” cells was calculated based on previously published transcriptomic analyses of adult MECs (Kendrick et al., 2008). The scores were computed using the Seurat function “AddModuleScore”.

#### 3D trajectory and pseudotime analysis

For this analysis, only the epithelial cell clusters from E13.5, E14.5, E15.5, and P0 were considered. The pre-processing steps previously described were re-applied (normalization, PCA, and basal and luminal score). Epithelial cells were then mapped in a 3D space including the luminal score and basal score on the x-axis and the PC related to developmental time on the y-axis. For each cell cluster, the coordinates of the center in the 3D space with the median for each dimension were calculated and called “pseudo-bulks”. A minimum spanning tree (MST) was generated to connect all pseudo-bulks. Basal and luminal trajectories were inferred through the MST.

To get the pseudotime of each cell along the basal or luminal trajectories, each cell was projected in the 3D space to the basal and luminal trajectories separately. Then, the pseudotime for each cell was defined as their distance from the initial point of the trajectory.

The luminal and basal gene expression heatmap was generated on the pseudotime with the “pheatmap” package. Briefly, the genes with the top 10% variation across cells within a lineage were selected. The gene expression values were smoothed versus the pseudotime using the generalized additive model (GAM). The hierarchical gene clusters were generated with Euclidean distance and Complete clustering algorithm.

#### Cell-cell interaction analysis

The cell-cell interaction analysis was performed using the CellPhoneDB version 3.0.0 (Vento-Tormo et al., 2018) with a p-value threshold of 0.01. The CellPhoneDB database is publicly available at https://www.cellphonedb.org/. It is a curated database of ligand-receptor interactions that allow to predict cell-cell interactions in transcriptomic data. CellPhoneDB was used on our scRNA-seq E15.5 dataset between both mesenchymal clusters (sub-epithelial cluster and dermal mesenchyme cluster) against the basal-like epithelial cell cluster.

### Statistics and Reproducibility

At least n=3 animals were used for each experiment, and experiments with at least n=3 replicates were used to calculate the statistical significance of each analysis. Statistical tests and further graphs were prepared in Prism (v9, GraphPad). All graphs show mean ± SEM. Differences between groups were assessed with two-tailed unpaired T-test with Welch’s correction. Statistical analyses between the localization of two RNA probes were assessed with two-way ANOVA test. The significance threshold was p < 0.05. * indicates p<0.05, ** indicates p<0.01, *** indicates p<0.001, and **** indicates p<0.0001.

## Data and software availability

Customized scripts and instructions are available from Github: https://github.com/Fre-Team-Curie/Embryo-mammary-gland.

The single cell RNA sequencing data have been deposited in NCBI’s Gene Expression Omnibus (GEO) repository and are accessible through GEO Series accession number GSE210594. All other data supporting the conclusions of this study are provided in the main text or the supplementary materials.

## Key Resource Table 1. Reagents and materials

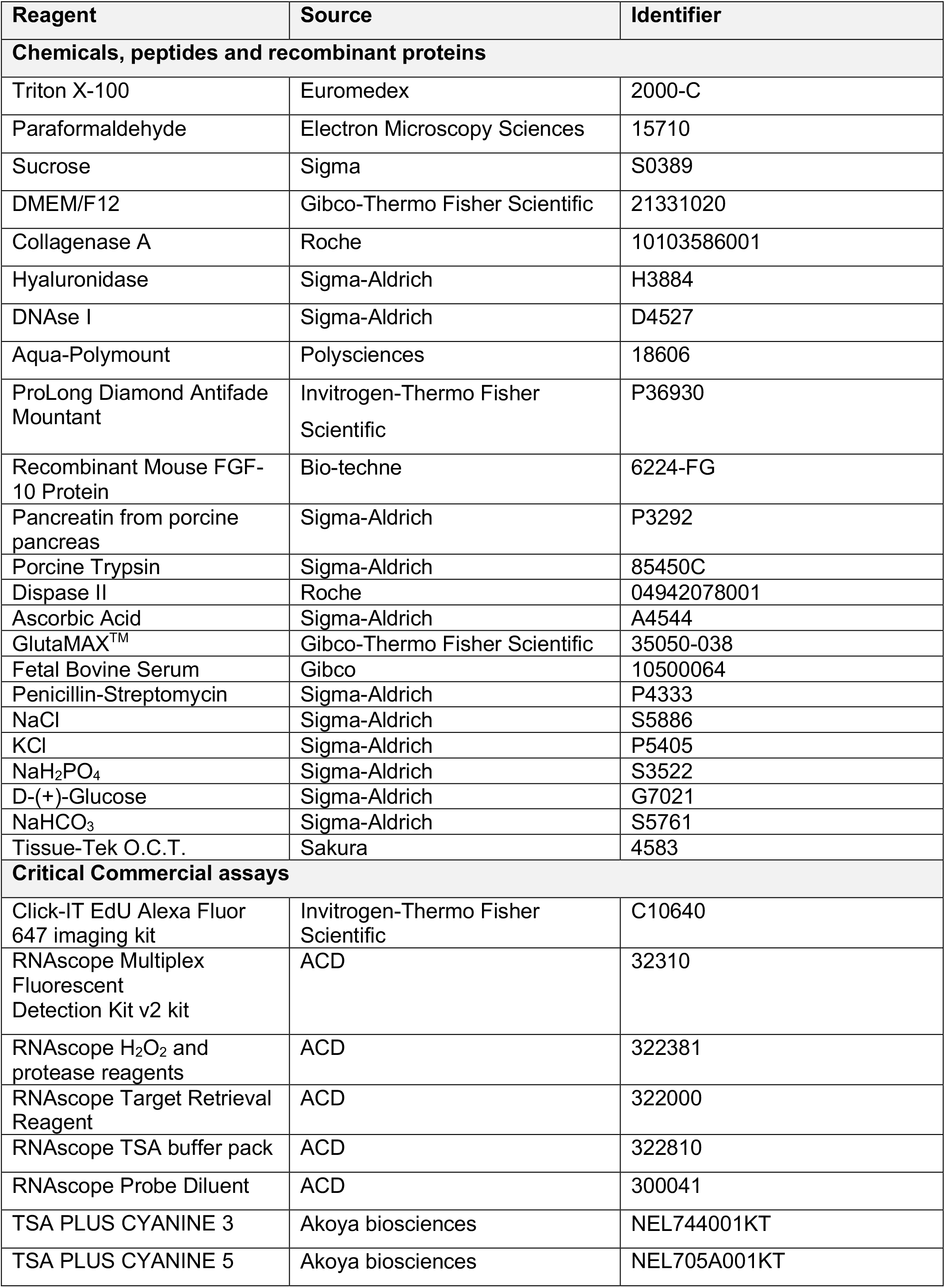
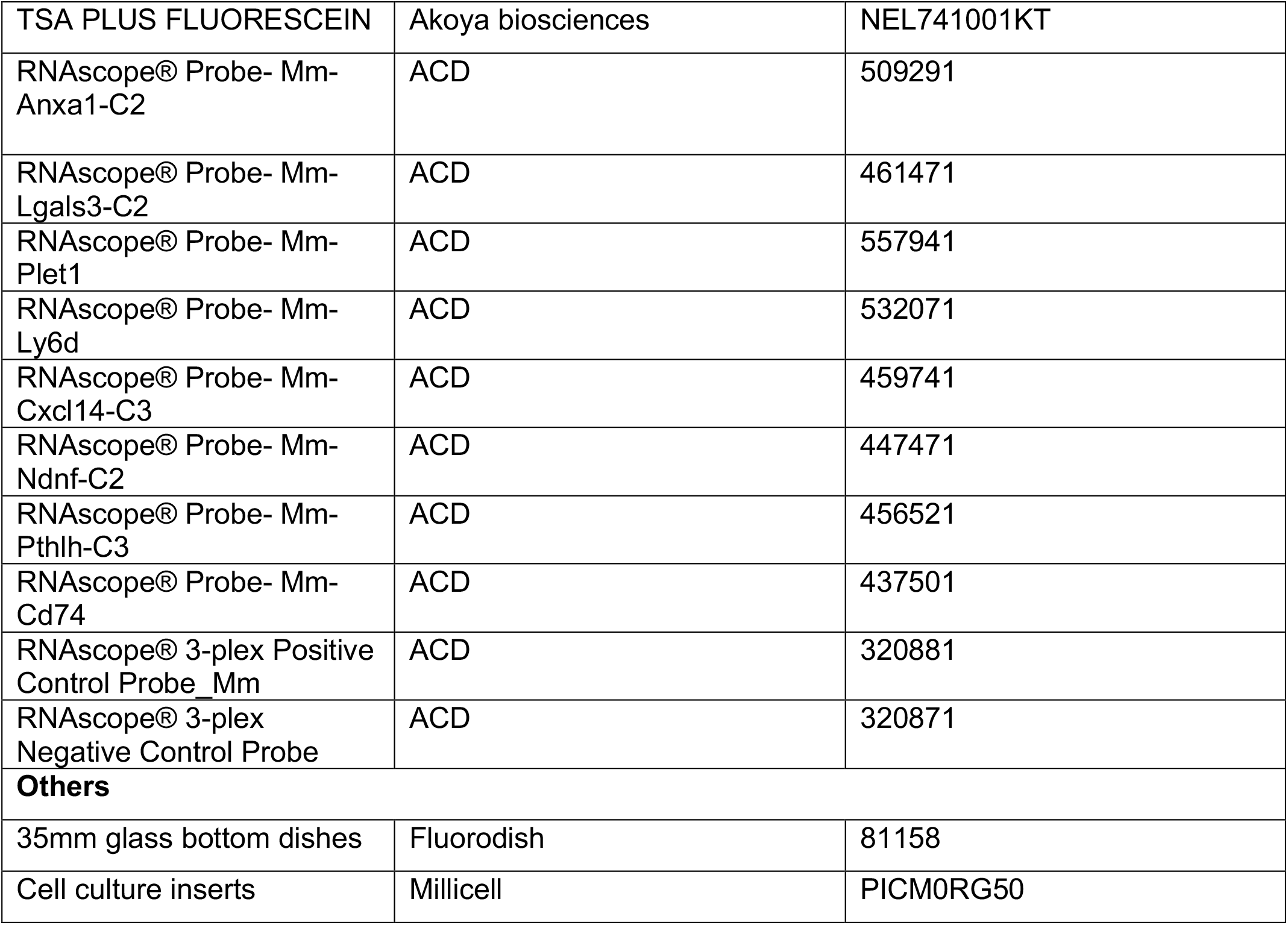

## Key Resource Table 2. Antibodies

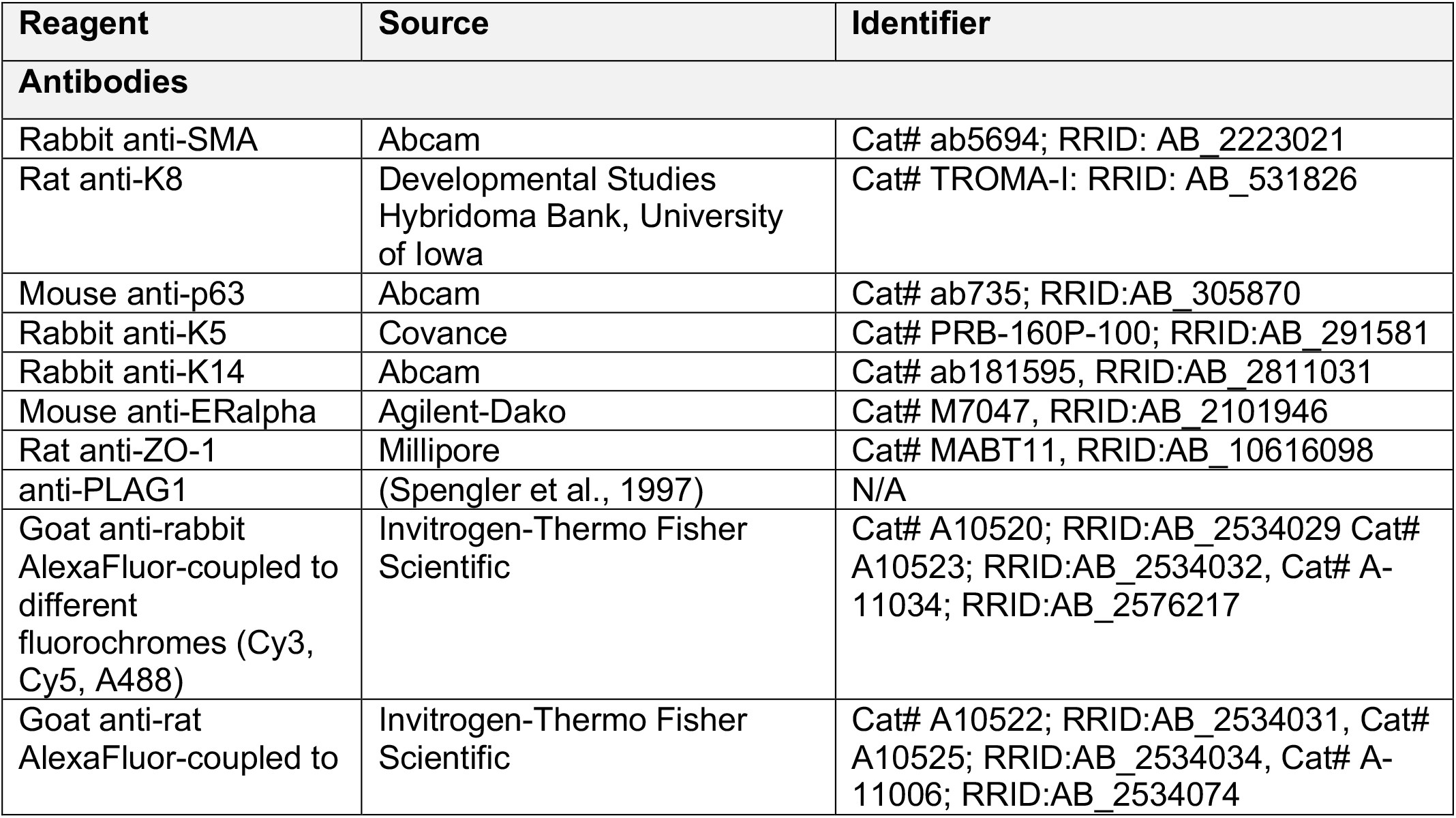
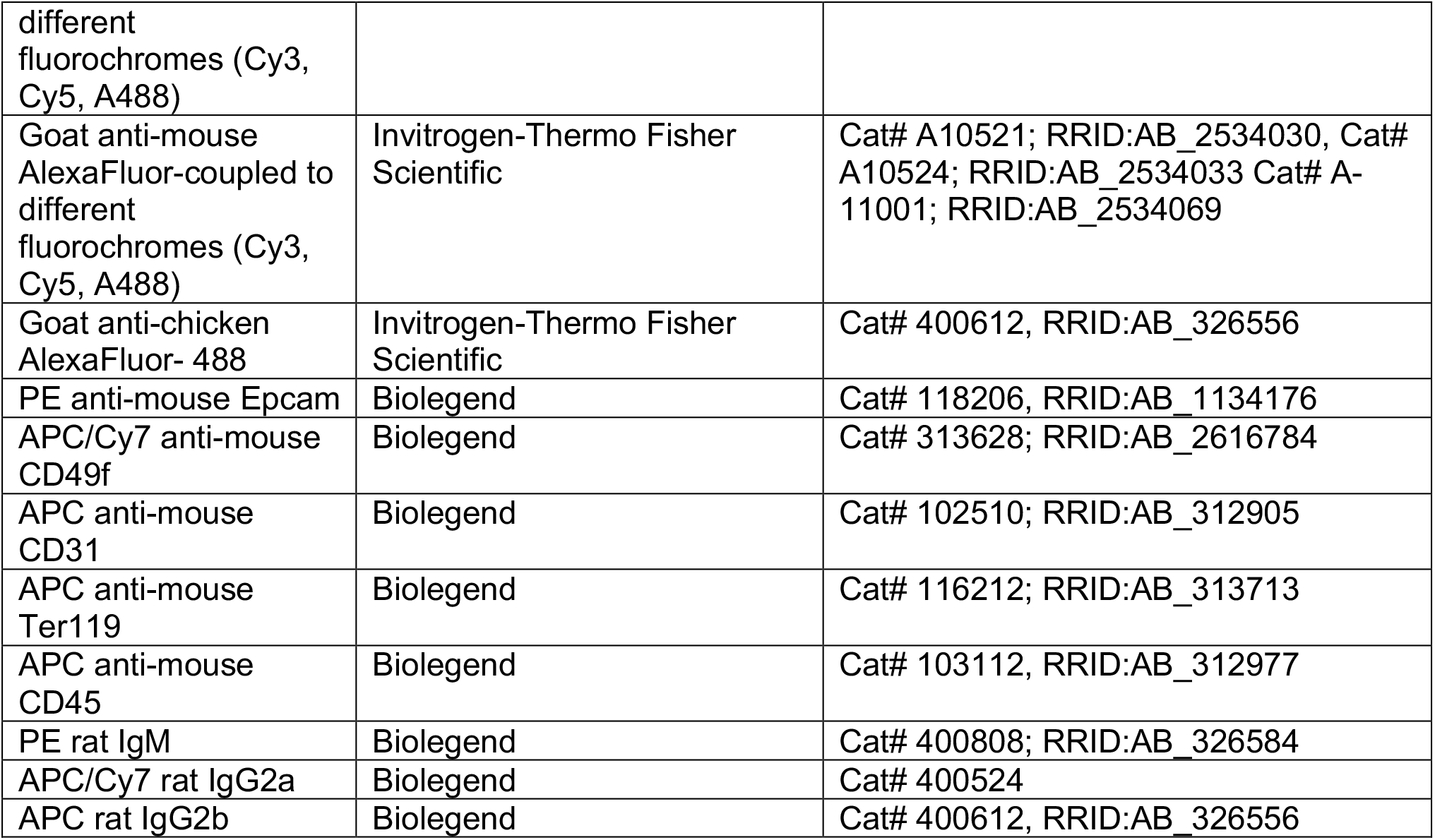

## Acknowledgements

The authors thank S. Tajbakhsh for the mTmG reporter line. We acknowledge the flow cytometry and cell sorting platform at Institute Curie for their technical support, the in vivo experimental facility for help in the maintenance and care of mouse colonies, the ICGex NGS platform of Institut Curie (supported by grants ANR-10-EQPX-03, Equipex and ANR-10-INBS-09-08, France Génomique Consortium, from the Agence Nationale de la Recherche “Investissements d’Avenir” program, by the Canceropole Ile-de-France and by the SiRIC-Curie program - SiRIC Grant INCa-DGOS-4654), the Cell and Tissue Imaging Platform-PICT-IBiSA (member of France-Bioimaging, ANR-10-INBS-04) of the Genetics and Developmental Biology Department (UMR3215/U934) for their expertise. The authors specially thank Olivier Leroy and Anne-Sophie Mace for image analysis support. We are grateful to members of the Fre laboratory for support and constructive discussions. This work was supported by Paris Sciences et Lettres (PSL* Research University) (grant # C19-64-2019-228), the French National Research Agency (ANR) grant numbers ANR-15-CE13-0013-01 and ANR-21-CE13-0047, the Medical Research Foundation FRM “FRM Equipes” EQU201903007821, the FSER (Fondation Schlumberger pour l’éducation et la recherche) FSER20200211117 and by Labex DEEP ANR-Number 11-LBX-0044 to S.F. C.C. was funded by the European Union’s Horizon 2020 research and innovation program under the Marie Skłodowska-Curie grant agreement No 666003 for PhD fellowships and the Medical Research Foundation FRM Project 12917. B.L.-L. is funded by a Vice-Chancellor’s Research Fellowship from the University of Bristol and acknowledges support from the Academy of Medical Sciences/Wellcome Trust/the Government Department of Business, Energy and Industrial Strategy/British Heart Foundation/Diabetes UK Springboard Award [SBF003/1170], Elizabeth Blackwell Institute for Health Research (University of Bristol), and Wellcome Trust Institutional Strategic Support Fund (204813/Z/16/Z).

The funders had no role in study design, data collection and analysis, decision to publish, or preparation of the manuscript.

## Author Contributions

C.C., B.L.-L., M.M.F., and S.F. conceived and designed the experiments. C.C., M.P. and M.M.F. performed all experiments. C.C., W.S. and R.J. performed the scRNA-seq data analysis. C.C., B.L.-L., V.K. and F.H. performed image analysis. C.C. and V.K. developed the image analysis pipeline. C.C., B.L.-L., M.M.F., and S.F. wrote the manuscript. All authors reviewed and approved the manuscript.

## References

Blaas, L., Pucci, F., Messal, H. A., Andersson, A. B., Ruiz, E. J., Gerling, M., Douagi, I., Spencer-Dene, B., Musch, A., Mitter, R., Bhaw, L., Stone, R., Bornhorst, D., Sesay, A. K., Jonkers, J., Stamp, G., Malanchi, I., Toftgard, R., & Behrens, A. (2016). Lgr6 labels a rare population of mammary gland progenitor cells that are able to originate luminal mammary tumours. Nature Cell Biology, 18(12), 1346–1356. https://doi.org/10.1038/ncb3434

Buechler, M. B., Pradhan, R. N., Krishnamurty, A. T., Cox, C., Calviello, A. K., Wang, A. W., Yang, Y. A., Tam, L., Caothien, R., Roose-Girma, M., Modrusan, Z., Arron, J. R., Bourgon, R., Müller, S., & Turley, S. J. (2021). Cross-tissue organization of the fibroblast lineage. Nature, 593(7860), 575–579. https://doi.org/10.1038/s41586-021-03549-5

Carabaña, C., & Lloyd-Lewis, B. (2022). Multidimensional Fluorescence Imaging of Embryonic and Postnatal Mammary Gland Development. In Vivanco MM (Ed.), Methods Mol Biol (Second, vol. 2471, pp. 19–48). https://doi.org/10.1007/978-1-0716-2193-6_2

Chan, C. J., Heisenberg, C. P., & Hiiragi, T. (2017). Coordination of Morphogenesis and Cell-Fate Specification in Development. Current Biology, 27(18), R1024–R1035. https://doi.org/10.1016/j.cub.2017.07.010

Chung, C.-Y., Ma, Z., Dravis, C., Preissl, S., Poirion, O., Luna, G., Hou, X., Giraddi, R. R., Ren, B., & Wahl, G. M. (2019). Single-Cell Chromatin Analysis of Mammary Gland Development Reveals Cell-State Transcriptional Regulators and Lineage Relationships. Cell Rep, 29(2), 495–510. https://doi.org/10.1016/j.celrep.2019.08.089

Davis, F. M., Lloyd-Lewis, B., Harris, O. B., Kozar, S., Winton, D. J., Muresan, L., & Watson, C. J. (2016). Single-cell lineage tracing in the mammary gland reveals stochastic clonal dispersion of stem/progenitor cell progeny. Nature Communications, 7(13053). https://doi.org/10.1038/ncomms13053

dos Santos, C. O., Rebbeck, C., Rozhkova, E., Valentine, A., Samuels, A., Kadiri, L. R., Osten, P., Harris, E. Y., Uren, P. J., Smith, A. D., & Hannon, G. J. (2013). Molecular hierarchy of mammary differentiation yields refined markers of mammary stem cells. Proceedings of the National Academy of Sciences of the United States of America, 110(18), 7123–7130. https://doi.org/10.1073/pnas.1303919110

Fankhaenel, M., Sadat Golestan Hashemi, F., Mosa Hosawi, M., Mourao, L., Skipp, P., Morin, X., LGJ Scheele, C., & Elias, S. (2021). Annexin A1 is a polarity cue that directs planar mitotic spindle orientation during mammalian epithelial morphogenesis. BioRxiv, 454117. https://doi.org/10.1101/2021.07.28.454117

Giraddi, R. R., Chung, C.-Y., Heinz, R. E., Balcioglu, O., Novotny, M., Trejo, C. L., Dravis, C., Hagos, B. M., Mehrabad, E. M., Rodewald, L. W., Hwang, J. Y., Fan, C., Lasken, R., Varley, K. E., Perou, C. M., Wahl, G. M., & Spike, B. T. (2018). Single-Cell Transcriptomes Distinguish Stem Cell State Changes and Lineage Specification Programs in Early Mammary Gland Development. Cell Rep, 24(6), 1653–1666. https://doi.org/10.1016/j.celrep.2018.07.025

Hafemeister, C., & Satija, R. (2019). Normalization and variance stabilization of single-cell RNA-seq data using regularized negative binomial regression. Genome Biology, 20(1). https://doi.org/10.1186/s13059-019-1874-1

Hannezo, E., Scheele, C. L. G. J., Moad, M., Drogo, N., Heer, R., Sampogna, Rosemary. V., van Rheenen, J., & Simons, B. D. (2017). A Unifying Theory of Branching Morphogenesis. Cell, 171(1), 242-255.e27. https://doi.org/10.1016/j.cell.2017.08.026

Hao, Y., Hao, S., Andersen-Nissen, E., Mauck, W. M., Zheng, S., Butler, A., Lee, M. J., Wilk, A. J., Darby, C., Zager, M., Hoffman, P., Stoeckius, M., Papalexi, E., Mimitou, E. P., Jain, J., Srivastava, A., Stuart, T., Fleming, L. M., Yeung, B., … Satija, R. (2021). Integrated analysis of multimodal single-cell data. Cell, 184(13), 3573-3587.e29. https://doi.org/10.1016/j.cell.2021.04.048

Huff, J. (2015). The Airyscan detector from ZEISS: confocal imaging with improved signal-to-noise ratio and super-resolution. Nature Methods, 12(12), i–ii. https://doi.org/10.1038/nmeth.f.388

Inman, J. L., Robertson, C., Mott, J. D., & Bissell, M. J. (2015). Mammary gland development: Cell fate specification, stem cells and the microenvironment. Development, 142(6), 1028–1042. https://doi.org/10.1242/dev.087643

Kendrick, H., Regan, J. L., Magnay, F. A., Grigoriadis, A., Mitsopoulos, C., Zvelebil, M., & Smalley, M. J. (2008). Transcriptome analysis of mammary epithelial subpopulations identifies novel determinants of lineage commitment and cell fate. BMC Genomics, 9, 591. https://doi.org/10.1186/1471-2164-9-591

Lan, Q., & Mikkola, M. L. (2020). Protocol: Adeno-Associated Virus-Mediated Gene Transfer in Ex Vivo Cultured Embryonic Mammary Gland. Journal of Mammary Gland Biology and Neoplasia, 25(4), 409–416. https://doi.org/10.1007/s10911-020-09461-4

Lilja, A. M., Rodilla, V., Huyghe, M., Hannezo, E., Landragin, C., Renaud, O., Leroy, O., Rulands, S., Simons, B. D., & Fre, S. (2018). Clonal analysis of Notch1-expressing cells reveals the existence of unipotent stem cells that retain long-term plasticity in the embryonic mammary gland. Nature Cell Biology, 20(6), 677–687. https://doi.org/10.1038/s41556-018-0108-1

Linkert, M., Rueden, C. T., Allan, C., Burel, J. M., Moore, W., Patterson, A., Loranger, B., Moore, J., Neves, C., MacDonald, D., Tarkowska, A., Sticco, C., Hill, E., Rossner, M., Eliceiri, K. W., & Swedlow, J. R. (2010). Metadata matters: Access to image data in the real world. Journal of Cell Biology, 189(5), 777–782. https://doi.org/10.1083/jcb.201004104

Lloyd-Lewis, B., Davis, F. M., Harris, O. B., Hitchcock, J. R., & Watson, C. J. (2018). Neutral lineage tracing of proliferative embryonic and adultmammary stem/progenitor cells. Development (Cambridge), 145(14). https://doi.org/10.1242/dev.164079

Mailleux, A., Spencer-Dene, B., Dillon, C., Ndiaye, D., Savona-Baron, C., Itoh, N., Kato, S., Dickson, C., Thiery, J., & Bellusci, S. (2002). Role of FGF10/FGFR2b signaling during mammary gland development in the mouse embryo. Development, 129(1), 53–60. https://doi.org/10.1242/dev.129.1.53

McInnes, L., Healy, J., & Melville, J. (2018). UMAP: Uniform Manifold Approximation and Projection for Dimension Reduction. http://arxiv.org/abs/1802.03426

Merrick, D., Sakers, A., Irgebay, Z., Okada, C., Calvert, C., Morley, M. P., Percec, I., & Seale, P. (2019). Identification of a mesenchymal progenitor cell hierarchy in adipose tissue. Science, 364(6438). https://doi.org/10.1126/science.aav2501

Miller, D. J., & Fort, P. E. (2018). Heat shock proteins regulatory role in neurodevelopment. Frontiers in Neuroscience, 12(821). https://doi.org/10.3389/fnins.2018.00821

Muzumdar, M. D., Tasic, B., Miyamichi, K., Li, N., & Luo, L. (2007). A global double-fluorescent cre reporter mouse. Genesis, 45(9), 593–605. https://doi.org/10.1002/dvg.20335

Pal, B., Chen, Y., Milevskiy, M. J. G., Vaillant, F., Prokopuk, L., Dawson, C. A., Capaldo, B. D., Song, X., Jackling, F., Timpson, P., Lindeman, G. J., Smyth, G. K., & Visvader, J. E. (2021). Single cell transcriptome atlas of mouse mammary epithelial cells across development. Breast Cancer Research, 23(1). https://doi.org/10.1186/s13058-021-01445-4

Prater, M. D., Petit, V., Alasdair Russell, I., Giraddi, R. R., Shehata, M., Menon, S., Schulte, R., Kalajzic, I., Rath, N., Olson, M. F., Metzger, D., Faraldo, M. M., Deugnier, M. A., Glukhova, M. A., & Stingl, J. (2014). Mammary stem cells have myoepithelial cell properties. Nature Cell Biology, 16(10), 942–950. https://doi.org/10.1038/ncb3025

Ronneberger, O., Fischer, P., & Brox, T. (2015). U-Net: Convolutional Networks for Biomedical Image Segmentation. http://arxiv.org/abs/1505.04597

Scheele, C. L. G. J., Hannezo, E., Muraro, M. J., Zomer, A., Langedijk, N. S. M., van Oudenaarden, A., Simons, B. D., & van Rheenen, J. (2017). Identity and dynamics of mammary stem cells during branching morphogenesis. Nature, 542(7641), 313–317. https://doi.org/10.1038/nature21046

Schmidt, U., Weigert, M., Broaddus, C., & Myers, G. (2018). Cell Detection with Star-convex Polygons. International Conference on Medical Image Computing and Computer-Assisted Intervention (MICCAI). https://doi.org/10.1007/978-3-030-00934-2_30

Sjöberg, E., Augsten, M., Bergh, J., Jirström, K., & Östman, A. (2016). Expression of the chemokine CXCL14 in the tumour stroma is an independent marker of survival in breast cancer. British Journal of Cancer, 114(10), 1117–1124. https://doi.org/10.1038/bjc.2016.104

Spengler, D., Villalba, M., Hoffmann, A., Pantaloni, C., Houssami, S., Bockaert, J., & Journot, L. (1997). Regulation of apoptosis and cell cycle arrest by Zac1, a novel zinc finger protein expressed in the pituitary gland and the brain. The EMBO Journal, 16(10), 2814–2825. https://doi.org/10.1093/emboj/16.10.2814

Spina, E., & Cowin, P. (2021). Embryonic mammary gland development. Seminars in Cell and Developmental Biology, 114, 83–92. https://doi.org/10.1016/j.semcdb.2020.12.012

Stuart, T., Butler, A., Hoffman, P., Hafemeister, C., Papalexi, E., Mauck, W. M., Hao, Y., Stoeckius, M., Smibert, P., & Satija, R. (2019). Comprehensive Integration of Single-Cell Data. Cell, 177(7), 1888-1902.e21. https://doi.org/10.1016/j.cell.2019.05.031

Tsang, S. M., Kim, H., Oliemuller, E., Newman, R., Boateng, N. A., Guppy, N., & Howard, B. A. (2021). Sox11 regulates mammary tumour-initiating and metastatic capacity in Brca1-deficient mouse mammary tumour cells. DMM Disease Models and Mechanisms, 14(5). https://doi.org/10.1242/DMM.046037

van Amerongen, R., Bowman, A. N., & Nusse, R. (2012). Developmental stage and time dictate the fate of Wnt/β-catenin-responsive stem cells in the mammary gland. Cell Stem Cell, 11(3), 387–400. https://doi.org/10.1016/j.stem.2012.05.023

van Keymeulen, A., Rocha, A. S., Ousset, M., Beck, B., Bouvencourt, G., Rock, J., Sharma, N., Dekoninck, S., & Blanpain, C. (2011). Distinct stem cells contribute to mammary gland development and maintenance. Nature, 479(7372), 189–193. https://doi.org/10.1038/nature10573

Vento-Tormo, R., Efremova, M., Botting, R. A., Turco, M. Y., Vento-Tormo, M., Meyer, K. B., Park, J. E., Stephenson, E., Polański, K., Goncalves, A., Gardner, L., Holmqvist, S., Henriksson, J., Zou, A., Sharkey, A. M., Millar, B., Innes, B., Wood, L., Wilbrey-Clark, A., … Teichmann, S. A. (2018). Single-cell reconstruction of the early maternal–fetal interface in humans. Nature, 563(7731), 347–353. https://doi.org/10.1038/s41586-018-0698-6

Voutilainen, M., Lindfors, P. H., & Mikkola, M. L. (2013). Protocol: Ex vivo culture of mouse embryonic mammary buds. Journal of Mammary Gland Biology and Neoplasia, 18(2), 239–245. https://doi.org/10.1007/s10911-013-9288-2

Wansbury, O., Mackay, A., Kogata, N., Mitsopoulos, C., Kendrick, H., Davidson, K., Ruhrberg, C., Reis-Filho, J. S., Smalley, M. J., Zvelebil, M., & Howard, B. A. (2011). Transcriptome analysis of embryonic mammary cells reveals insights into mammary lineage establishment. Breast Cancer Research, 13(4). https://doi.org/10.1186/bcr2928

Watson, C. J., & Khaled, W. T. (2020). Mammary development in the embryo and adult: New insights into the journey of morphogenesis and commitment. Development, 147(22). https://doi.org/10.1242/dev.169862

Weigert, M., Schmidt, U., Haase, R., Sugawara, K., & Myers, G. (2020). Star-convex Polyhedra for 3D Object Detection and Segmentation in Microscopy. Proceedings of the IEEE/CVF Winter Conference on Applications of Computer Vision (WACV), 3666–3673. https://doi.org/10.48550/arXiv.1908.03636

Wuidart, A., Ousset, M., Rulands, S., Simons, B. D., van Keymeulen, A., & Blanpain, C. (2016). Quantitative lineage tracing strategies to resolve multipotency in tissue-specific stem cells. Genes Dev, 30(11), 1261–1277. https://doi.org/10.1101/gad.280057.116

Wuidart, A., Sifrim, A., Fioramonti, M., Matsumura, S., Brisebarre, A., Brown, D., Centonze, A., Dannau, A., Dubois, C., van Keymeulen, A., Voet, T., & Blanpain, C. (2018). Early lineage segregation of multipotent embryonic mammary gland progenitors. Nature Cell Biology, 20(6), 666–676. https://doi.org/10.1038/s41556-018-0095-2

