## Supplementary Figures for "Positional cues underlie cell fate specification during branching morphogenesis of the embryonic mammary epithelium"

Supplementary Figure 1. Lineage committed cells exist in early MG development.

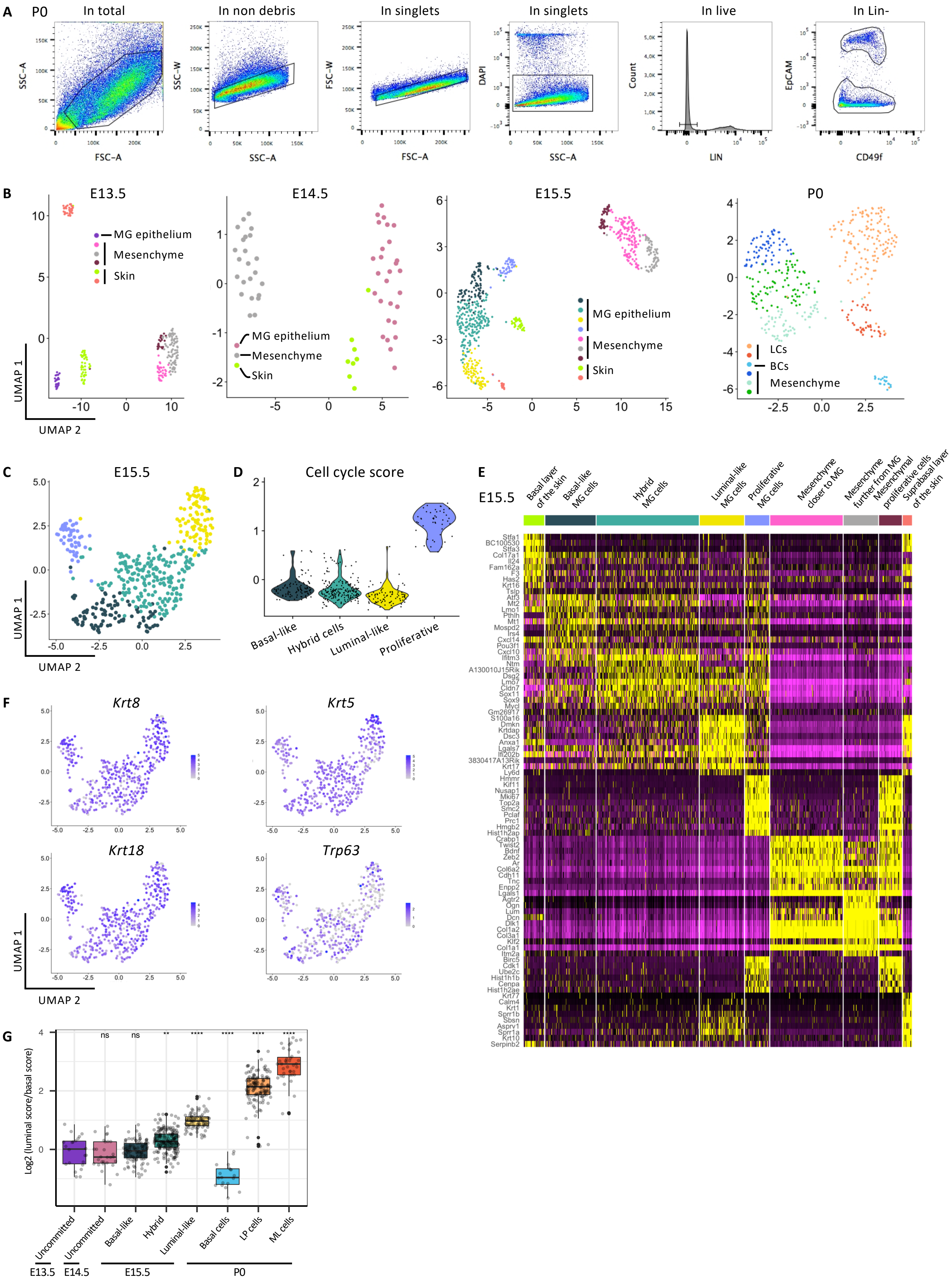

**Supplementary Figure 1. Related to Figure 1. Lineage committed cells exist in early MG development.**

(A) Representative FACS dot plots of the gating strategy used to sort P0 epithelial and mesenchymal cells. (B) UMAP plots of embryonic MECs and surrounding mesenchymal cells isolated by scRNA-seq at E13.5, E14.5, E15.5 and P0. Cells are color-coded by cluster. (C) UMAP plot of embryonic MECs isolated at E15.5 after subset analysis of all MECs (including proliferative cells shown in light blue). (D) Violin plot representation of the cell cycle score in each mammary epithelial cluster at E15.5. (E) Heatmap showing the expression of genes specific for each cell cluster at E15.5. Each column is color-coded according to the cell cluster from (B). (F) UMAP plots from (C) showing the expression of specific luminal (*Krt8* and *Krt18*) and basal (*Krt5* and *Trp63*) genes commonly used to distinguish adult LCs and BCs but unable to discriminate distinct cell clusters at E15.5. (G) Box plots illustrating the log2 fold change of the luminal/basal score ratio in each cluster. ** $p<0.01$, **** $p<0.0001$, ns: non-significant.

Supplementary Figure 2. Identification of novel genes that distinguish lineage-biased embryonic mammary cells.

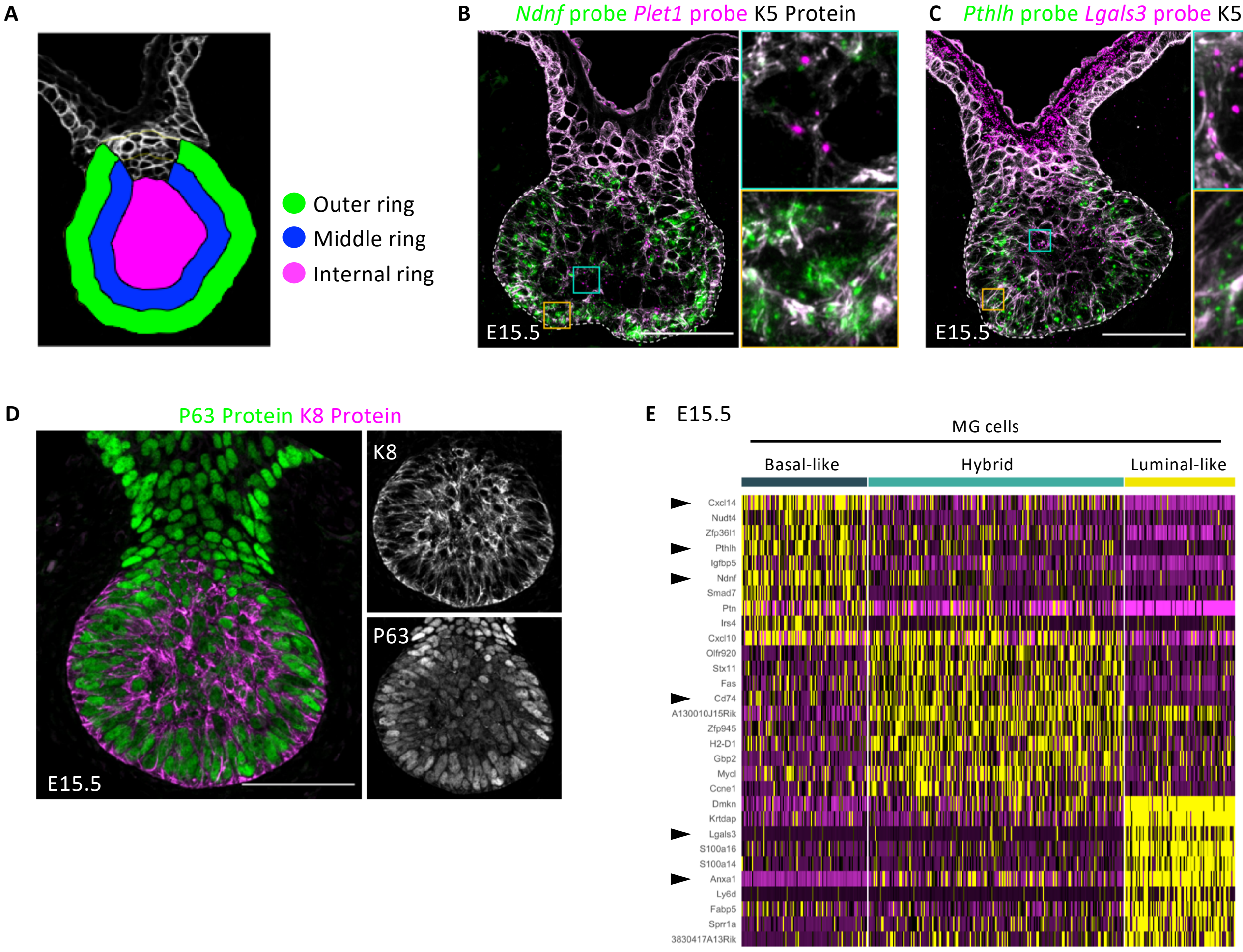

**Supplementary Figure 2. Related to Figure 3. Identification of novel genes that distinguish lineage-biased embryonic mammary cells.**

(A) Cartoon illustrating the outer ring (in green), middle ring (in blue) and internal ring (in magenta) used in our unbiased quantitative analysis. (B and C) Representative sections of embryonic mammary buds at E15.5 showing the expression of *Ndnf* (basal gene, in green) and *Plet1* (luminal gene, in magenta) (B) or *Pthlh* (basal gene, in green) and *Lgals3* (luminal gene, in magenta) (C), detected by RNAscope and immunostained with K5 (in white). (D) Single optical section showing the expression of the luminal epithelial marker K8 (in magenta), and the basal epithelial marker P63 (in green) in an embryonic mammary bud at E15.5. K8 and P63 are co-expressed by all MECs at E15.5. (E) Heatmap illustrating the expression of genes specific for each MEC cluster at E15.5. Each column is color-coded according to the cell cluster from Figure 1B. Black arrowheads indicate genes used in RNAscope experiments.

Supplementary Figure 3. The heterogeneity of mesenchymal cells increases at birth.

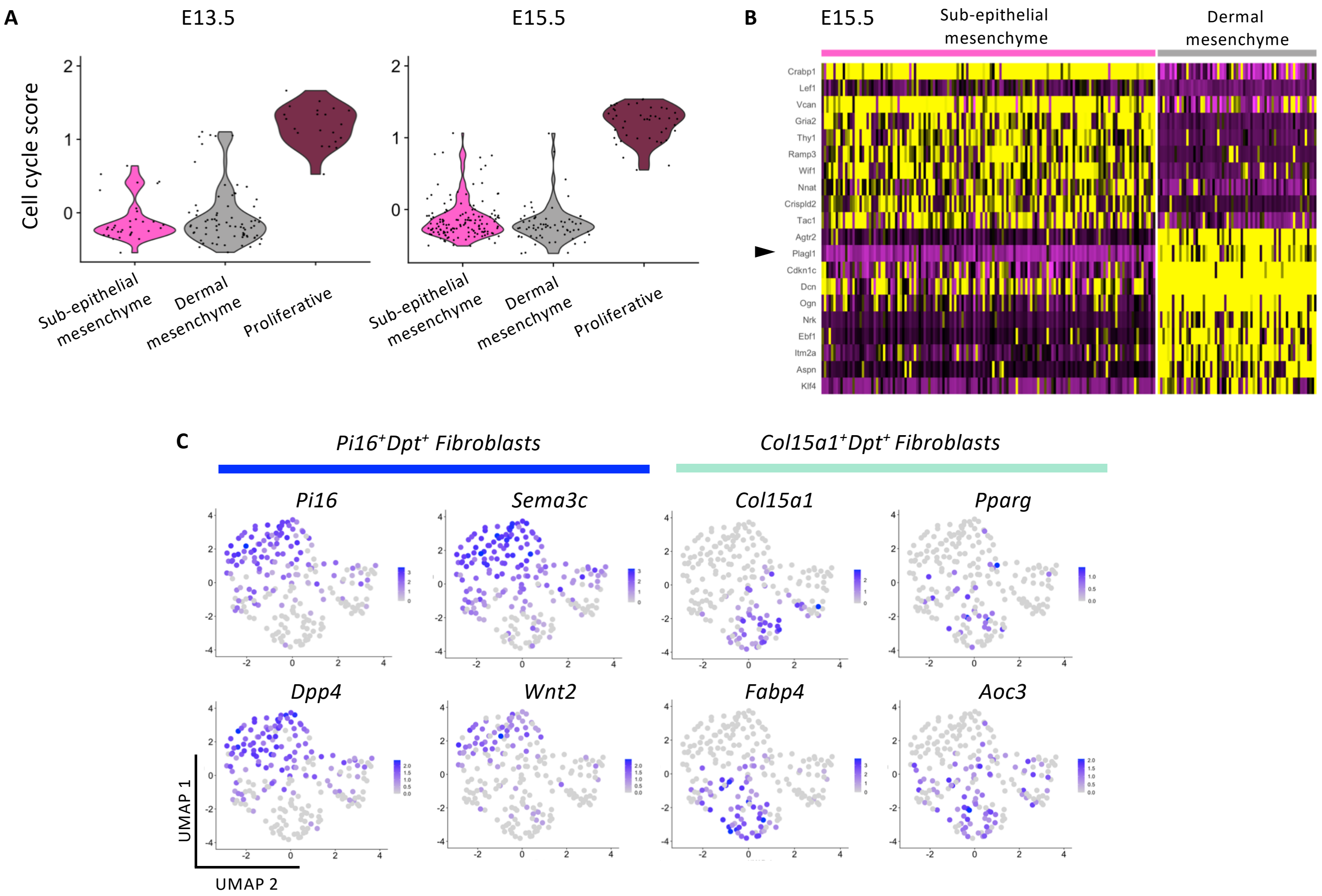

**Supplementary Figure 3. Related to Figure 4. The heterogeneity of mesenchymal cells increases at birth.**

(A) Violin plots representing the cell cycle score in each mammary mesenchymal cluster at E13.5 and E15.5. (B) Heatmap illustrating the expression of genes specific for each mesenchymal cluster at E15.5. Each column is color-coded according to the cell cluster from Figure 4A. The black arrowhead indicates *Plagl1*, previously used to label the dermal mesenchyme. (C) UMAP plots from Figure 4A illustrating the expression of cluster-specific genes in mesenchymal cells at P0.

Supplementary Figure 4. Ligand-receptor interaction pairs between both mesenchymal populations and the basal-like cluster at E15.5.

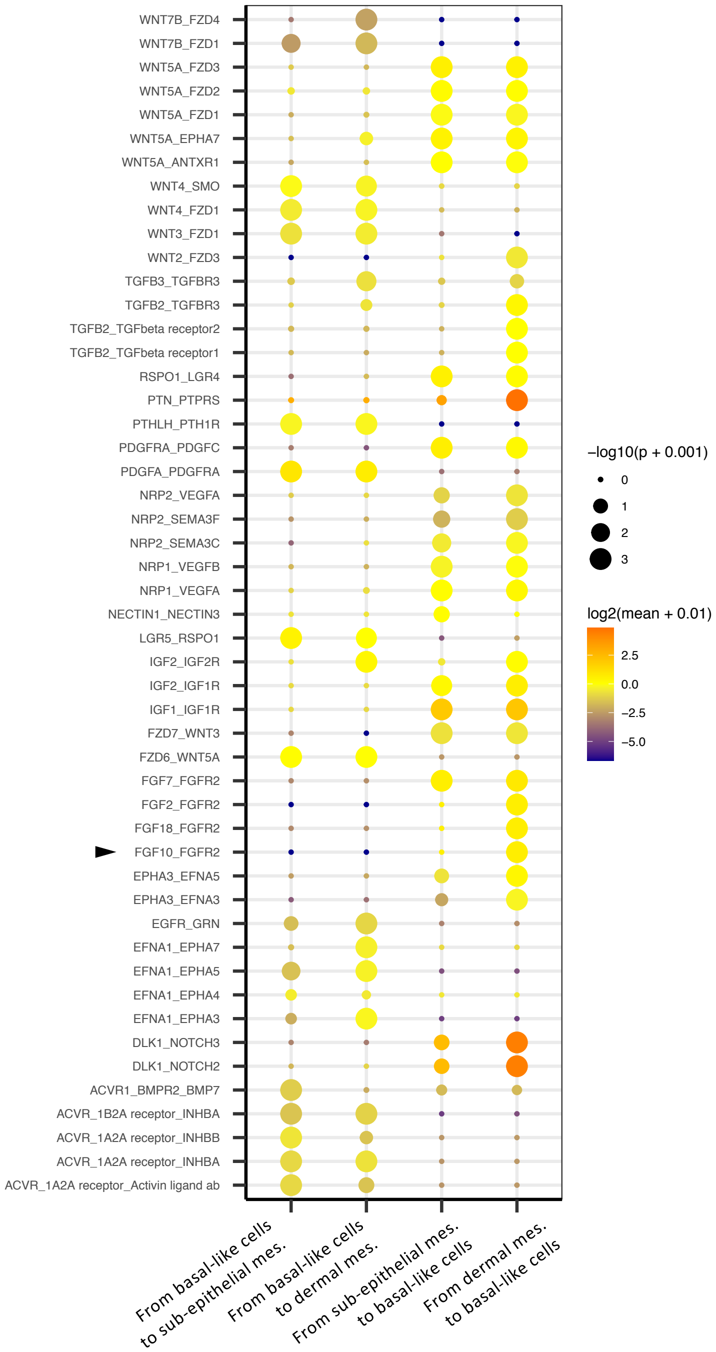

**Supplementary Figure 4. Related to Figure 5. Ligand-receptor interaction pairs identified between the two mesenchymal populations and the basal-like cluster at E15.5.** CellPhoneDB analysis with the predicted ligand-receptor interactions between the two mesenchymal populations, sub-epithelial or dermal mesenchyme, and basal-like cells at E15.5 and vice versa ($p$-value < 0.01). The arrowhead highlights the ligand-receptor interaction between FGF10 and FGFR2 that was functionally investigated in embryonic *ex vivo* cultures.

Supplementary Figure 5. Mammary bud *ex vivo* cultures recapitulate embryonic mammary morphogenesis and epithelial lineage segregation.

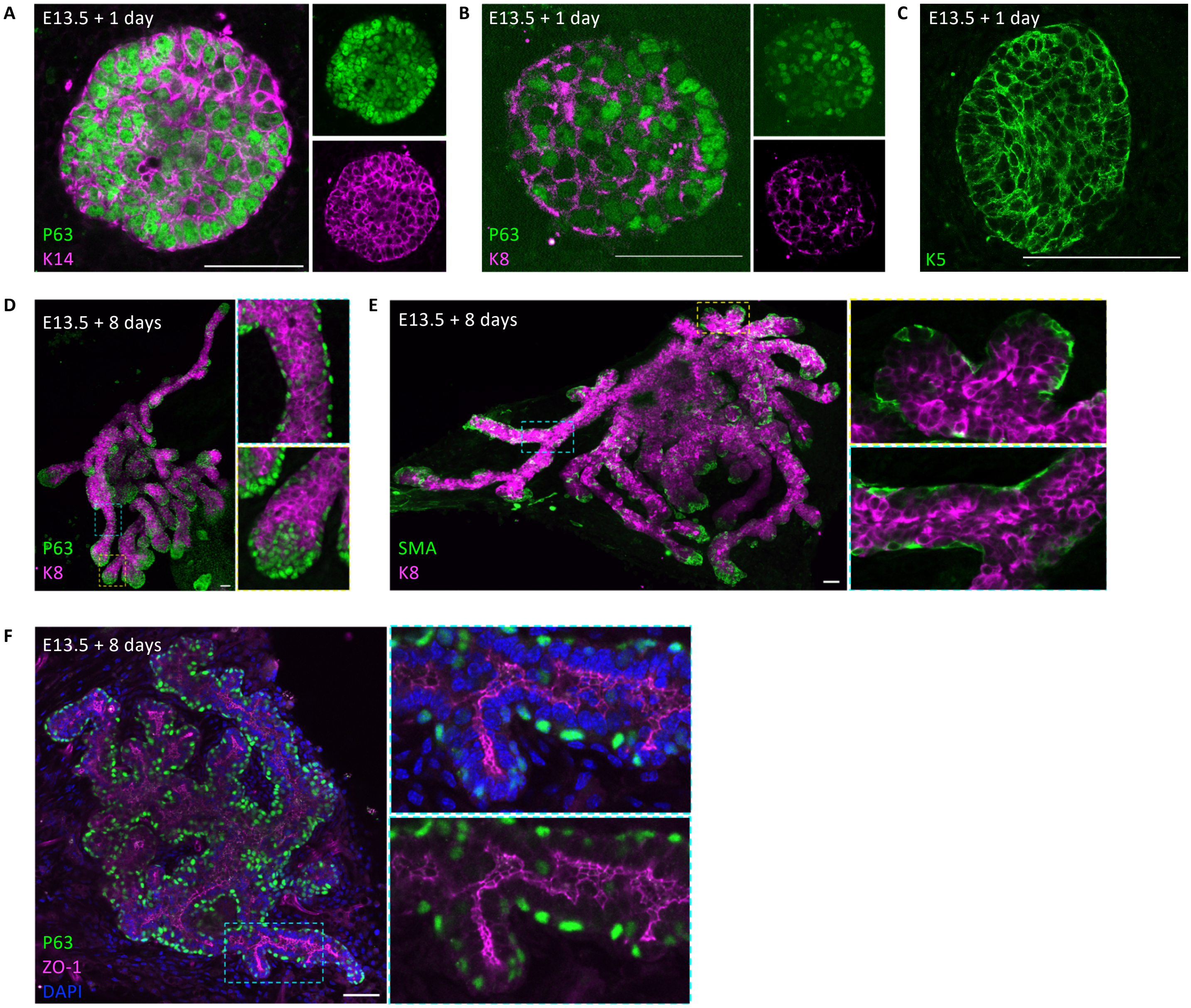

**Supplementary Figure 5. Related to Figure 5. Mammary bud *ex vivo* cultures recapitulate embryonic mammary morphogenesis and epithelial lineage segregation.**

(A-C) Representative images of mammary embryonic buds dissected at day E13.5 and cultured *ex vivo* for 1 day, immunostained for the following lineage markers: P63 (in green) and K14 (in magenta) (A), P63 (in green) and K8 (in magenta) (B), and K5 (in green) (C). (D-F) Representative images of mammary embryonic buds dissected at day E13.5 and cultured *ex vivo* for 8 days, immunostained for the following lineage and polarity markers: P63 (in green) and K8 (in magenta) (D), $\alpha$-SMA (in green) and K8 (in magenta) (E), and P63 (in green) and ZO-1 (in magenta) (F). Scale bars: 50 $\mu$m (in A, B and C), 100 $\mu$m (in D, E and F).
